# Functional insights into a peculiar tetra-modular LPMO from the human pathogen *Enterobacter cloacae*

**DOI:** 10.1101/2025.07.04.663164

**Authors:** Koyel Bardhan, Saumashish Mukherjee, Theruvothu Madathil Vandhana, Alessia Munzone, Andres Posbeyikian, Sacha Grisel, Lal Duhsaki, Jean-Guy Berrin, Bastien Bissaro, Jogi Madhuprakash

**Affiliations:** Department of Plant Sciences, School of Life Sciences, University of Hyderabad, Gachibowli, Hyderabad-500046, Telangana, India; INRAE, Aix Marseille University, UMR1163 Biodiversité et Biotechnologie Fongiques, 13009 Marseille, France; The Sainsbury Laboratory, University of East Anglia, Norwich Research Park, NR4 7UH, Norwich, UK

**Keywords:** *Enterobacter*, Chitin, LPMO, AA10, CBM73, Synergy, Copper coordination

## Abstract

*Enterobacter cloacae* is a Gram-negative nosocomial human pathogen that inhabits diverse ecological niches. Its genome encodes a conserved set of putative chitin-active enzymes, including a peculiar lytic polysaccharide monooxygenase (LPMO), termed *Ec*LPMO, which we functionally characterized in this study. *Ec*LPMO is a tetra-modular protein consisting of an auxiliary activity family 10 (AA10) catalytic domain, two central domains of unknown function (DUF-A and DUF-B), and a C-terminal carbohydrate-binding module (CBM73). Functional assays using full-length *Ec*LPMO and its truncated variants demonstrated that the AA10 domain oxidatively cleaves chitin at the C1 position. The CBM73 module enhances chitin binding and promotes synergy with endogenous chitinases. Notably, *Ec*LPMO displayed a particularly strong synergistic effect with the unimodular chitinase *Ec*ChiA, leading to up to 14-fold and 60-fold increases in GlcNAc release from α- and β-chitin, respectively. Deletion of both DUFs reduced *Ec*LPMO activity. While DUF-A alone and the association of DUF-A and DUF-B showed limited chitin binding, DUF-B alone exhibited no binding, suggesting a distinct role. Unexpectedly, using state-of-the-art structural modelling (AlphaFold3), we observed that the DUF-B domain contains two highly conserved histidines that coordinate the AA10-bound copper, forming a previously unreported ‘inter-domain tetra-histidine copper coordination’ center. These findings highlight the structural and functional complexity of *Ec*LPMO and suggest that its accessory domains, particularly DUF-B, may contribute to enzyme stability and substrate interaction. We speculate that DUF-B may protect the LPMO active site from oxidative damage, a feature that could prove crucial in its ecological and pathogenic contexts.

## 1. Introduction

Lytic polysaccharide monooxygenases (LPMOs) are copper-dependent enzymes that mediate oxidative cleavage of glycosidic bonds in crystalline polysaccharides such as chitin (1) and cellulose (2-4). This oxidative mechanism introduces nicks on the polysaccharide surface, disrupting its structure and facilitating more efficient hydrolysis by glycoside hydrolases (GHs) (5, 6). At their active site, LPMOs contain a mononuclear copper ion coordinated by two conserved histidine residues, known as the ‘histidine brace’ (7). To become catalytically active, the copper must first be reduced from Cu(II) to Cu(I), typically by external electron donors such as small organic compounds (e.g., ascorbate) or enzymatic partners (3, 8-10). Once reduced, and in the presence of hydrogen peroxide, the LPMO-Cu(I) can then catalyze hydroxylation of either the C1 or C4 carbon of the scissile glycosidic bond, triggering cleavage via a peroxygenase-type mechanism (11-15). The resulting products are typically aldonic acids (C1-oxidation) or geminal diols (C4-oxidation).

LPMOs are currently grouped in the Carbohydrate-Active enZymes (CAZy) database as Auxiliary Activities (AA), spanning eight families (AA9–11 and AA13–17) (16). Chitin-active LPMOs are primarily found in families AA10 (1), AA11 (17), and AA15 (18). While AA11s are restricted to fungi, AA15 LPMOs occur across viruses, algae, oomycetes, and insects. The AA10 family stands out as the most taxonomically diverse, with representatives from bacteria (1), viruses (19, 20), plants (21), and, as more recently unveiled, fungi (22). Of note, most of these AA10 LPMOs characterized to date act on chitin.

Similar to other CAZymes, many LPMOs are multi-modular, incorporating additional domains such as carbohydrate-binding modules (CBMs), GH domains, or glycosylphosphatidylinositol (GPI) anchors. These domains can modulate LPMO binding affinity, catalytic efficiency, and stability (23). They are typically connected by flexible linkers that influence interdomain interactions and substrate accessibility (24). In addition, many LPMOs contain disordered C-terminal regions (dCTR), which are present across most LPMO families (except AA13), and still remain largely functionally uncharacterized (25). Regarding AA10 LPMOs, to date, approximately 23 multi-modular LPMOs have been experimentally characterized for activity on chitin or cellulose.

Beyond their recognized role in biomass degradation, LPMOs are now being implicated in diverse biological functions (26), including pathogenesis and cellular development (27-31). Notably, several bacterial AA10 LPMOs have been linked to virulence. One early example is GbpA from *Vibrio cholerae*, a chitin-active LPMO involved in mucin binding and intestinal colonization (32). More recently, CbpD from *Pseudomonas aeruginosa* was identified as a key virulence factor supporting bacterial survival in the human bloodstream (27).

In the course of identifying LPMO candidates from human-associated bacterial pathogens, we focused on the Gram-negative nosocomial pathogen *Enterobacter cloacae. E. cloacae* is a clinically relevant opportunistic pathogen associated with hospital-acquired infections, particularly in immunocompromised individuals. It can cause respiratory, urinary tract infections, and meningitis (33). Due to its increasing antibiotic resistance, it is classified among the WHO-designated ESKAPE pathogens (34). Genome analysis revealed a conserved, single-copy, multi-modular AA10 LPMO across 46 complete *E. cloacae* genomes (NCBI; **Table S1**), which we refer to as *Ec*LPMO. Its modular organization shows resemblance to previously studied virulence-associated LPMOs such as GbpA (from *V. cholerae*; 32), CbpD (from *P. aeruginosa;* 27) and *Vca*LPMO10A (from *V. campbellii*; 35), despite relatively low sequence identity. This conservation and structural similarity suggest that *Ec*LPMO may play a biologically important role, potentially linked to the host-pathogen interface.

In this study, we aimed to characterize the chitinolytic machinery of *E. cloacae*, with a specific focus on *Ec*LPMO. Using sequence and structural *in silico* analyses, we predicted functional properties and designed truncated protein variants. We then assessed the ability of *Ec*LPMO and its variants to bind and oxidize crystalline chitin, providing insight into the role of its accessory domains. Finally, we examined the synergistic effects of *Ec*LPMO with *E. cloacae* chitinases on α-and β-chitin, advancing our understanding of how this enzyme may contribute to efficient chitin degradation and potentially to pathogenicity.

## 2. Materials and methods

### 2.1. Bioinformatic analysis

The annotation of chitin-active CAZymes for different strains of *E. cloacae* (including the one under study, i.e., ATCC 13047^T^) was conducted using the dbCAN3 meta-server (36). An integrated approach using all three designated tools/databases of the dbCAN3 server i.e., HMMER:dbCAN, Diamond:CAZy and HMMER:dbCAN-sub, was employed for CAZyme prediction and annotation. Only those proteins or domains annotated by at least two tools were considered for further analysis (36). We also present the frequency of strains that possess ‘n’ members of a given CAZy family, illustrating the distribution of chitin-active CAZymes. (**Table S2**).

Domain architecture of *Ec*LPMO and few other chitin-active CAZymes (*Ec*ChiA, *Ec*ChiB, *Ec*ChiC and *Ec*HexA) from *E. cloacae* ATCC 13047^T^ was predicted using Pfam (https://pfam.xfam.org/) and the NCBI Conserved Domain Database https://www.ncbi.nlm.nih.gov/Structure/cdd/cdd.shtml). SignalP 5.0 (http://www.cbs.dtu.dk/services/SignalP/) was used to predict the presence of signal peptide. Genetic arrangement and genomic localization of these CAZymes in the representative type strain genome, *E. cloacae* ATCC 13047^T^ was identified using the RAST servers.

The modularity of *Ec*LPMO was further analyzed by the expert annotation of Prof. B. Henrissat (CAZy database, Marseille, France), who determined the boundaries of the different domains (i.e. AA10, DUF-A, DUF-B and CBM73). Sequence alignment of *Ec*LPMO with its homologs was performed using the T-Coffee server (https://tcoffee.crg.eu/) and the output was generated using ESPript 3.0.

### 2.2. Phylogenetic analysis

To build the phylogenetic tree of AA10s, we used 176 AA10 sequences including sequences belonging to clades with characterized members, and sequences belonging to clades still uncharacterized. Of note, the N-terminal signal peptide and the variable C-terminal regions of all AA10 sequences were cut manually using BioEdit (37) in order to keep the catalytic domain only. The non-redundancy of sequences was verified using CD-HIT. The 176 AA10 sequences were aligned using MAFFT-DASH (L-INS-i method) (38), which includes structural data input. The resulting multiple sequence alignments were used to infer the phylogenetic trees via the IQ-Tree webserver (39), with the standard bootstrap analysis (100 iterations) and a SH-aLRT branch test (1,000 replicates).

To build the phylogenetic tree of CBM73s, 5,025 sequence IDs of CBM73-containing proteins were retrieved from the CAZy database, which were used to retrieve the sequences from the NCBI database. The sequences were aligned with MAFFT-FFT-large-NS-2 (38) and, upon inspection of (predicted/experimental) structures, the region of about 60 amino acids corresponding to the CBM73 (40) was extracted from the MSA using BioEdit software. To decrease the number of sequences in order to build a reliable phylogenetic tree, CBM73 proteins sharing a sequence identity > 90% were clustered using CD-HIT (41), resulting in 572 clusters. A representative sequence of each cluster (provided by the CD-HIT software) was used to build the phylogenetic tree. This pool of sequences includes sequences of characterized CBM73, *i.e.*, the CBM73s from *Pseudomonas aeruginosa* PA01 (GenBank ID AAG04241.1, UniProt ID Q91589) (42), from *Cellvibrio japonicus* Ueda107 (GenBank ID ACE83992.1, UniProt ID B3PJ79, (NMR) PDB ID 6Z41) (43) and from *Vibrio campbellii* ATCC BAA-1116 (GenBank ID ABU72648.1, PDB ID 8GUM) (35). To build the tree, the 572 sequences were aligned with MAFFT v7 and the resulting multiple sequence alignment was used to infer the phylogenetic trees via the IQ-Tree webserver (39), with the ultrafast bootstrap analysis (1,000 iterations) and a SH-aLRT branch test (1,000 replicates). Each branch was then assigned with SH-aLRT and UFBoot supports, allowing to reliably define clades when SH-aLRT ≥ 80% and UFBoot ≥ 95%. Sequences forming each clade were retrieved and re-aligned with MAFFT v7 to generate a sequence conservation plot using WebLogo 3.0 (44). The phylogenetic trees were visualized in iTOL (45) and edited in Illustrator CC 2017.

### 2.3. Structure determination using AlphaFold

Mature wild-type *Ec*LPMO and *Vc*GbpA were modelled using AlphaFold3 via the AlphaFold Server (46). Corresponding histidine-to-alanine variants (*Ec*LPMO^H354A,H384A^ and *Vc*GbpA^H357A,H388A^) were also modelled. For both wild-type and mutant proteins, structural predictions were generated either in the apo form or in complex with Cu(II). Default template mode was used, with the PDB template cutoff set to 30/09/2021, and model generation was repeated across five fixed seed values (Seed = 1–5).

A custom bash script was used to extract confidence metrics from the AlphaFold server output, including pTM, ipTM, and PAE plots, and to generate box plots of ipTM scores across the five seed replicates. Structural models were visualized using UCSF ChimeraX (47).

A custom Python script was written to identify protein atoms located within 3.6 Å of the modelled copper ion, extracting all atomic contacts for each predicted structure. This data was used to generate UpSet plots in Python, summarizing how many seed replicates (out of 5) predicted contacts between Cu(II) and the four conserved histidine residues: His22, His109, His354, and His384 for *Ec*LPMO, and His24, His121, His357, and His388 for *Vc*GbpA.

### 2.4. Cloning, expression and purification of recombinant enzymes

The full-length LPMO gene, *Ec*LPMO^FL^ (residues 1-484) including the native signal peptide (residues 1-21) was amplified from the genomic DNA of *E. cloacae* ATCC 13047 using the primers listed in **Table S3.** Both the insert and pET-22b(+) expression vector were double digested using the restriction endonucleases *Nde*I and *Xho*I (Thermo Scientific, USA), and were ligated using the Rapid Ligation kit (ThermoScientific, USA). The entire ligation mixture was then transformed by heat-shock into chemically competent *E. coli* DH5α cells. Positive clones were screened using colony PCR and the sequence integrity was further confirmed by Sanger sequencing. The same strategy was followed for generating the truncations of *Ec*LPMO^FL^, which were, *Ec*LPMO^ΔCBM73^ (residues 1-413), *Ec*LPMO^CD^ (residues 1-190), the DUFA-DUFB double domains (residues 205-413), and the single domain DUF-A (residues 205-308), DUF-B (residues 316-413) and CBM73 (residues 424-484) using the primer combinations as listed in the **Table S3.**

Sequence-validated clones were transformed by heat-shock into chemically competent BL21 (DE3) pLysS *E. coli* cells. Colonies obtained were inoculated into lysogenic broth (LB) containing 100 μg/mL ampicillin. *Ec*LPMO^FL^, *Ec*LPMO^ΔCBM73^, DUF-A and CBM73 were expressed by growing the transformants at 37°C for 3 h (until the OD reached 0.4-0.6) followed by induction with 0.4 mM IPTG for 16 h at 18°C and shaking at 180 rpm. A similar strategy was followed for the double domain DUFA-DUFB and the single domain DUF-B, but induction was performed with 0.7 mM IPTG. *Ec*LPMO^CD^ was expressed by growing at 30°C for 16 h without IPTG induction. The cells were harvested by centrifugation at 8000 rpm for 15 min. All the AA10-containing variants, i.e. *Ec*LPMO^FL^, *Ec*LPMO^ΔCBM73^ and *Ec*LPMO^CD^, all contained the native signal peptide and were thus isolated from the periplasmic fraction (SP cleaved off upon export) using an osmotic shock method essentially as described earlier by Madhuprakash et al., (2015) (48). The resulting periplasmic fractions were dialyzed against the equilibration buffer (50 mM NaH_2_PO_4_, 300 mM NaCl and 10 mM imidazole, pH 8.0), prior to purification. The truncated variants, i.e. DUFA-DUFB, DUF-A, DUF-B and CBM73, were produced intracellularly and thus isolated by sonication of the cells, as follows: the cell pellets were resuspended in equilibration buffer and disrupted by pulsed sonication for 5 min on ice (cycles of 20 s on and 40 s off), and the cell debris were removed by centrifugation at 18,000 rpm for 45 min at 4°C. The supernatant obtained was filtered through a 0.45 µm membrane filters (Merck Millipore, USA) and purification was performed by Ni-NTA affinity chromatography as reported previously (49). Fractions with the highest purity were pooled together and concentrated by centrifugation at 3,200 rpm using Vivaspin® Turbo 15 Ultrafiltration Units of 10 kDa MWCO (Sartorius, Germany). The final concentrated protein fractions were subjected to buffer exchange against 50 mM sodium phosphate, pH 8.0. The protein concentration was estimated using Pierce BCA (Bicinchoninic acid) assay kit (Thermo Scientific, USA) as per the manufacturer’s protocol.

The GH18 chitinases, *Ec*ChiA (uni-modular) and *Ec*ChiB (multi-modular) were also cloned from the genomic DNA of *E. cloacae* ATCC 13047 into pET-28a(+) and pET-22b(+) expression vectors, respectively, under the restriction sites, *Nco*I and *Xho*I (Thermo Scientific, USA). Sequence-validated clones were transformed by heat-shock into chemically competent BL21 (DE3) pLysS *E. coli* cells. Colonies obtained were inoculated into lysogenic broth (LB) containing 50 μg/mL kanamycin (in case of *Ec*ChiA) or 100 μg/mL ampicillin (in case of *Ec*ChiB). The recombinant chitinases were expressed by growing the transformants at 37°C for 3 h (until the OD reached 0.4-0.6) followed by induction with 0.4 mM IPTG for 16 h at 18°C and shaking at 180 rpm. The cells were harvested by centrifugation at 8000 rpm for 15 min. The cell pellet were resuspended in equilibration buffer containing 0.1 mM phenylmethylsulfonyl fluoride and sonicated using a Sonic-Ultrasonic Processor VCX-750 sonicator. The sonication was performed at 30% amplitude for 5 min at 20 s on and 40 s off cycles. Frequent monitoring of the sample was done to avoid heating. The sonicated sample was then centrifuged at 18000 rpm for 45 min at 4°C to separate the pellet and supernatant. The supernatant was purified by Ni-NTA affinity chromatography, followed by concentration and buffer exchange using Vivaspin® Turbo 15 Ultrafiltration Units of 10 kDa MWCO (Sartorius, Germany) as mentioned above. Protein estimation was done using Pierce BCA (Bicinchoninic acid) assay kit (Thermo Scientific, USA) as per the manufacturer’s protocol.

### 2.5. LPMO reaction and products analysis

#### 2.5.1. MALDI-TOF mass spectrometric analysis of oxidized products

The reaction mixture was composed of 1 μM of *Ec*LPMO^FL^, 10 mg/ml of α-or β -chitin (Stellar Biosol, Gujarat, India), 1 mM ascorbate, in 5 mM Tris-Cl pH 8.0. The reaction was incubated at 37°C for 24 h, at 1,000 rpm in an Eppendorf Thermomixer® C (Eppendorf India Pvt. Ltd., Chennai, India). Samples were collected at 12 h and 24 h and filtered using a 96-well filter plate (0.45 μm filters; Merck Millipore, USA) operated by a Millipore vacuum manifold. The filtered samples were lyophilized and analysed by MALDI-TOF-MS analysis as described previously (1).

#### 2.5.2. Detection of H_2_O_2_ production by Amplex red assay

The apparent hydrogen peroxide generation during substrate-free *Ec*LPMO reactions was followed by using the Amplex Red/HRP assay as established by Kittl et al. (2012) (50). Briefly, a pre-reaction mixture contained (final concentrations after all additions): HRP (0.1 mg/mL), Amplex red (200 µM) and *Ec*LPMO^FL^, *Ec*LPMO^CD^, *Ec*LPMO^ΔCBM73^ or Apo-*Ec*LPMO^ΔCBM73^ (1 µM) in sodium phosphate buffer (pH 7.0, 50 mM). The final reaction (100 µL in a 96-well microplate), conducted at 25°C, was initiated by the addition of ascorbate (50 µM). The formation of resorufin (2:1 ratio with respect of H_2_O_2_), was monitored at 575 nm over a 90-minute duration using a microplate reader (BioTek Epoch 2, Agilent technologies, US). The background of non-enzymatic H_2_O_2_ production was assessed under identical conditions and no LPMO and was comparable to the reaction with the non-metalated LPMO (“Apo”). H_2_O_2_ solutions, prepared in TraceSELECT water were used to generate a standard curve under the same conditions. The concentration of H_2_O_2_ stock solution was verified at 240 nm (ε_240_ = 43.6 M^-1^.cm^-1^).

#### 2.5.3. Product analysis using HPAEC-PAD

Time course degradation experiments were performed by mixing (in a total reaction volume of 1 mL) 1 μM of enzyme (*Ec*LPMO^FL^, *Ec*LPMO^CD^ or *Ec*LPMO^ΔCBM73^) and 10 mg/mL of α-or β-chitin in 50 mM Tris-Cl pH 8.0. The reaction was initiated by addition of 1 mM ascorbate. The reaction was performed at 37°C and 1000 rpm for 24 h in an Eppendorf ThermoMixer C®. An aliquot of 150 μL was collected at different time points and filtered using a 96-well filter plate operated by a Millipore vacuum manifold and the filtrate was analyzed using high performance anion exchange chromatography with pulsed amperometric detection (HPAEC-PAD, DIONEX ICS6000 system, Thermo Fisher Scientific, Waltham, MA, USA), as previously described (22). The system was equipped with a CarboPac-PA1 guard column (2 x 50 mm) and a CarboPac-PA1 column (2 x 250 mm) kept at 30°C. All reactions were performed in triplicate with appropriate control reactions under the similar experimental conditions as required.

### 2.6. Binding assay on chitin substrates

The binding assays were conducted following the protocol previously described by Forsberg et al. (2016) (40) with slight modifications. Each reaction contained 10 mg/mL of α-or β-chitin substrates with 100 μg/mL of protein in 50 mM sodium phosphate buffer pH 7.0. The reaction was incubated for different time points (2.5, 5, 7.5, 10, 15, 30, 45 and 60 minutes) using an Eppendorf Thermomixer C® (Eppendorf India Pvt. Ltd., Chennai, India) at 1000 rpm. These reactions were performed at 22°C for CBM73 and at 4°C for the other truncations i.e., *Ec*LPMO^FL^, *Ec*LPMO^CD^ and *Ec*LPMO^ΔCBM73^. These temperatures were standardized based on the quality of binding isotherms observed for each truncated variant.

Binding studies for the independent *Ec*LPMO domains, i.e., DUF-A+B, DUF-A and DUF-B on both α-and β-chitin were performed at 22°C at different time points (0, 6, 12, 24, 48 h for DUF-A+B double domain; 0, 15, 30, 45, 60, 90, 120, 180 min for DUF-A and DUF-B single domains). Binding affinities of these truncations were verified under different pH conditions, i.e., 50 mM sodium phosphate pH 8.0, 50 mM bis-tris pH 6.0 and 50 mM CAPS pH 10.0, keeping all other reaction parameters constant.

Binding kinetics were performed to obtain the equilibrium binding constants (*K*_d_) and substrate binding capacity (*B*_max_) by using different concentrations of the protein (0, 10, 20, 50, 75, 100, 150, and 300 μg/mL) in 50 mM sodium phosphate pH 7.0 with 10 mg/mL α-or β-chitin. Firstly, a standard curve of the individual proteins was prepared in 50 mM sodium phosphate pH 7.0 without the substrate. In all conditions, i.e. in the presence and absence of the substrate, the reaction mixtures were incubated at 22°C for CBM73 and at 4°C in case of *Ec*LPMO^FL^, *Ec*LPMO^CD^ and *Ec*LPMO^ΔCBM73^ for 2 h in an Eppendorf Thermomixer C® at 1,000 rpm. Post-incubation, the reaction mixtures were centrifuged at 11,000 rpm for 10 min at 4°C. The quantity of protein in the supernatant (referred to as “free protein”, P_free_) was analyzed by the Pierce Bicinchoninic Acid (BCA) assay kit (Thermo Scientific, USA). All the assays were performed in triplicates with suitable controls. *K*_d_ and *B*_max_ were determined by fitting the binding isotherms to the one-site binding equation: [*P*_bound_] = *B*_max_ [*P*_free_]/*K*_d_ +[*P*_free_], where *P* represents the protein, by non-linear regression using the GraphPad Prism software (GraphPad Software Inc., San Diego, CA).

### 2.7. Synergy experiments for chitin degradation

Synergy experiments were performed by incubating 10 mg/mL of α-or β-chitin with 1 μM of *Ec*LPMO^FL^ or its truncated variants (*Ec*LPMO^CD^ and *Ec*LPMO^ΔCBM73^), in combination with 1 μM of *Ec*ChiA or *Ec*ChiB, in 50 mM Tris-Cl pH 8.0, in the presence of 1 mM ascorbate. All the reactions were performed in triplicate, and incubated at 40°C, 1,000 rpm in an Eppendorf ThermoMixer C®, up to 24 h. Aliquots were collected at different time points and filtered using a 96-well filter plate (0.45 μm filters; Merck Millipore) operated by a Millipore vacuum manifold. The filtrates were mixed with equal volume of 70% acetonitrile and analyzed by HPLC on the Shim-pack GIST NH2 column (4.6 mm ID X 250 mm, Shimadzu, Japan) through isocratic elution using 70% acetonitrile at a flow rate of 0.7 mL/min. CHOS were detected by monitoring the absorbance at 210 nm. Quantification of the neutral chitooligosaccharides (CHOS) generated was performed essentially as described previously (51, 52).

## 3. Results

### 3.1. Bioinformatic analyses of EcLPMO

To get preliminary insights into *Ec*LPMO properties and function, we carried out sequence and phylogenetic analyses. *Ec*LPMO^FL^ is a multi-modular LPMO comprising a total of 484 amino acids. It includes a 21-residue N-terminal signal peptide, followed by an AA10 catalytic domain (residues 22-190), two domains of unknown function DUF-A (residues 205-308) and DUF-B (residues 316-413) and a C-terminal CBM73 domain (residues 424-484) (**Fig. 1A and B)**. A phylogenetic analysis of the catalytic domain only of 176 AA10 LPMOs showed that *Ec*AA10 belongs to a clade of well-known chitin-active AA10s (**Fig. 1C**). In details, *Ec*AA10 shares a sequence identity of 51.5%, 48.8%, and 33.5% with the AA10 catalytic domains of GbpA (PDB: 2XWX), *Vca*LPMO10A (PDB: 8GUM), and CbpD (PDB: 7SQX), respectively, which are all chitin-active ‘GbpA-type’ LPMOs (27, 32, 35). Multiple sequence alignment revealed the conservation of the ‘histidine brace’ residues (His1 and His88 in *Ec*LPMO mature sequence (i.e. without SP); **Fig. S1**).

**Fig. 1.**
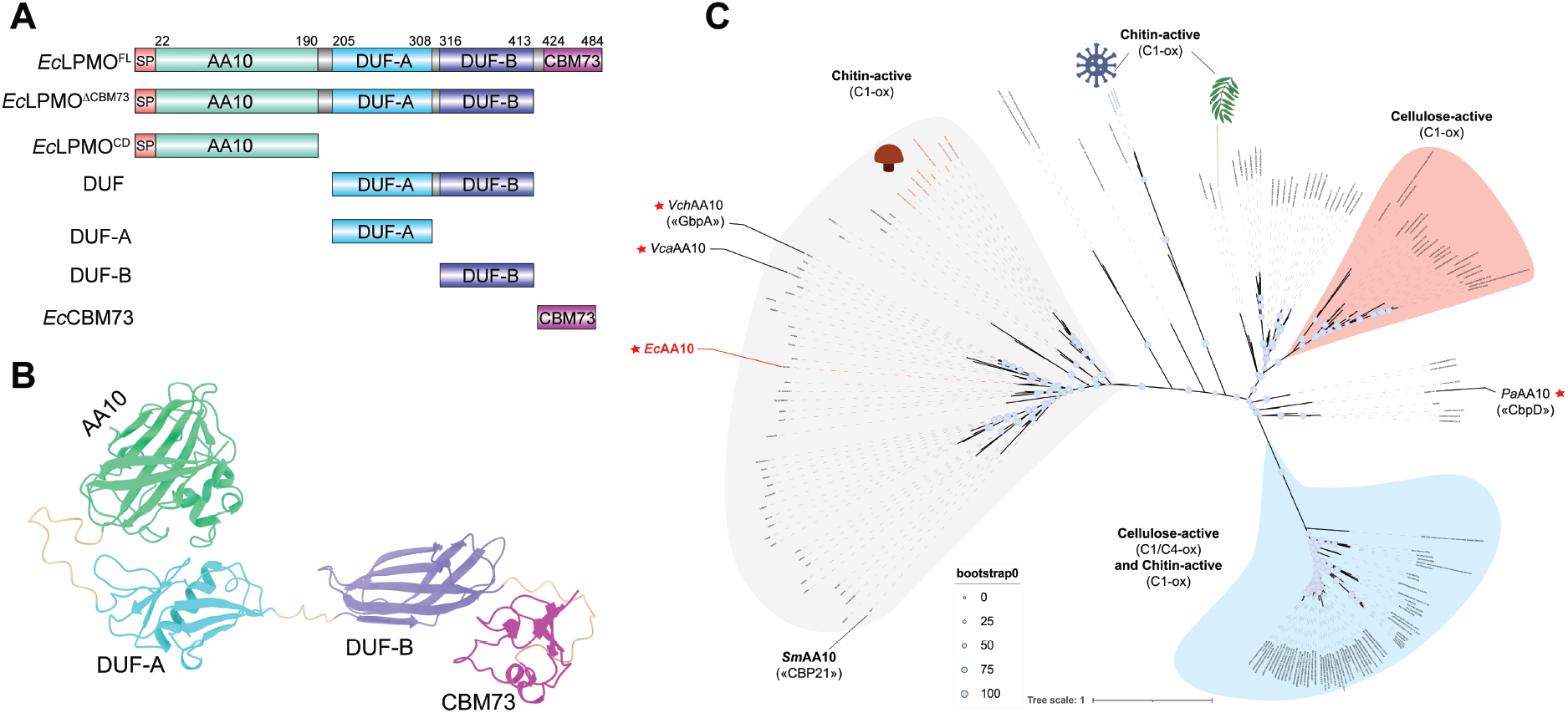
Sequence analysis of *Ec*LPMO. (A) Domain architecture of full-length *Ec*LPMO (*Ec*LPMO^FL^) and its truncated variants. The constructs, including *Ec*LPMO^FL^, *Ec*LPMO^ΔCBM73^, and *Ec*LPMO^CD^, contain a predicted signal peptide (SP; residues 1–22). (B) AlphaFold3-predicted structure of *Ec*LPMO^FL^, highlighting its modular domain organization. (C) Phylogenetic tree of the catalytic domains from 176 AA10 LPMOs. Bootstrap values are shown at branch nodes as blue circles (see scale in figure). Red filled stars mark previously characterized tetra-modular LPMOs orthologs of *Ec*LPMO (printed in red).

Regarding the DUF domains, DUF-A shares sequence identities of 40%, 37.6% and 34% with the GbpA_D2_ (GbpA_2 domain), the *Vca*LPMO10A-GbpA_2 and CbpD-MX (GbpA_2 domain), respectively. Meanwhile, DUF-B has a relatively low sequence identity of 30.1% and 24.7% compared to GbpA_D3_ and *Vca*LPMO10A-MX, respectively.

To evaluate the evolutionary relationship of the CBM73 domain with other members of the CBM73 family, we performed a phylogenetic analysis using 572 representative sequences clustered at 90% identity from 5,025 CBM73 entries available in the CAZy database. The resulting phylogenetic tree revealed five major clades, which likely reflect underlying functional divergence within the family (**Fig. 2A**). Notably, the CBM73 family remains largely uncharacterized, with most clades lacking any biochemically or structurally studied members, likely due to its relatively recent identification (40). Interestingly, approximately half of the representative sequences fall within a single superclade (highlighted in purple), which can be further resolved into multiple subclades. *Ec*CBM73 clusters within this purple superclade, alongside two previously characterized CBM73s, *Pa*CBM73 (42) and *Vca*CBM73 (35), though each resides in a separate subclade. *Ec*CBM73 shares only moderate sequence identity with these known homologs: 33.3% with *Cj*CBM73, 49.2% with *Pa*CBM73, and 60.3% with *Vca*CBM73. Structural superimposition of *Ec*CBM73 model (AF3) with X-ray crystallographic structures of *Pa*CBM73 and *Vca*CBM73 shows high similarity (RMSD values of 0.298 Å and 0.45 Å for C^α^ atoms, respectively). In contrast, superposition with *Cj*CBM73 revealed more pronounced structural differences, with an RMSD of 1.508 Å, supporting the phylogenetic separation of these CBMs.

**Fig. 2:**
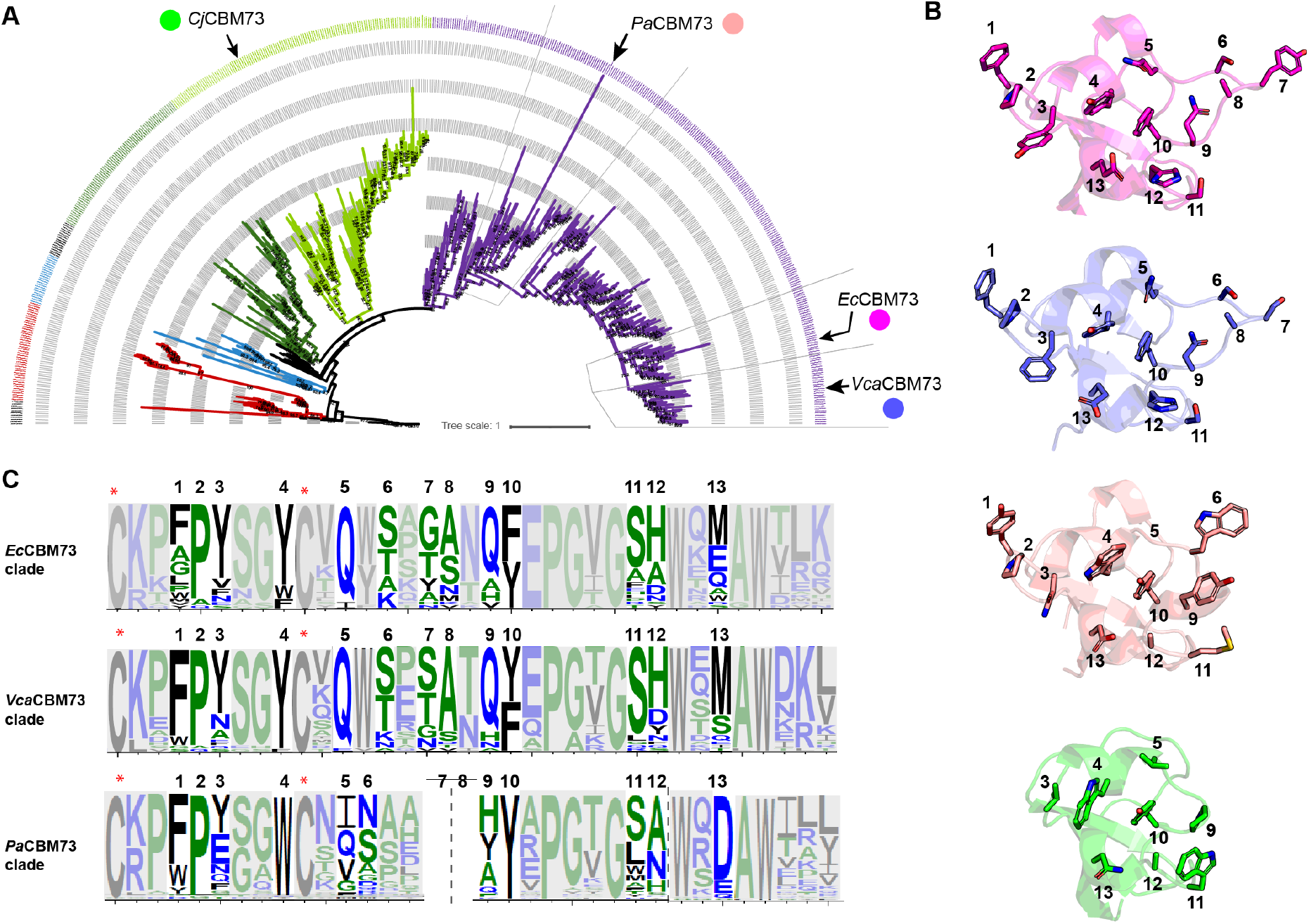
Phylogenetic tree of the CBM73 domain. **(A)** The tree was built with the CBM73 domain of 572 sequences, representative (on the basis of sequence clustering with 90% identity threshold) of 5,025 sequences annotated in the CAZy database. The tree was inferred using IQ-Tree (52) (1,000 ultrafast bootstraps; values displayed on the tree) and visualized with the Interactive Tree Of Life (iTOL) software (45). Characterized CBM73s, namely from *Pseudomonas aeruginosa* PA01 (GenBank ID AAG04241.1) (42), from *Cellvibrio japonicus* Ueda107 (GenBank ID ACE83992.1) (43), and from *Vibrio campbellii* ATCC BAA-1116 (GenBank ID ABU72648.1) (35) are highlighted on the tree. **(B)** Substrate-binding surface view highlighting solvent-exposed residues (shown as sticks) of, from top to bottom, *Ec*CBM73 (AF model), *Vca*CBM73 (PDB 8GUM; 35), *Pa*CBM73 (AF model) and *Cj*CBM73 (AF model). **(C)** WebLogo of the MSA of subclades (delimited by grey lines in panel A) containing *Ec*CBM73, *Pa*CBM73 and *Vca*CBM73, and corresponding to the stretch of amino acids forming the substrate binding surface (shown in panel B). Residues not highlighted in panel B are masked by a grey shading in panel C. The same numbering of highlighted residues has been used in both panels B and C. Note that some residues shown in panel B have a very low frequency of occurrence within the subclade (e.g. Trp at pos. 6 in *Pa*CBM73), and are thus barely readable in the corresponding WebLogo shown in panel C.

### 3.2. Comparison of CBM73 substrate-binding surfaces and sequence features

To further investigate the potential functional implications of these evolutionary relationships, the substrate-binding surfaces of *Ec*CBM73, *Vca*CBM73, *Pa*CBM73, and *Cj*CBM73 were compared based on their modelled structures (**Fig. 2B**). The three CBM73s from the purple clade (*Ec*CBM73, *Vca*CBM73, and *Pa*CBM73) exhibit more extended binding surfaces relative to *Cj*CBM73, which presents a compact and shallow architecture. Among the extended forms, *Ec*CBM73 displays the broadest surface (∼31.1 Å), followed by *Vca*CBM73 (∼28.7 Å) and *Pa*CBM73 (∼21.3 Å), while *Cj*CBM73 exhibits the narrowest (∼14.2 Å). These differences likely reflect clade-specific substrate recognition strategies and may point to distinctions in binding affinity.

Sequence logos derived from each CBM73 subclade (**Fig. 2C**) reveal partial conservation at key positions, such as a highly conserved proline at position 2, likely important for maintaining the core structural framework. However, other regions exhibit notable divergence. For instance, comparing *Ec*CBM73 to *Vca*CBM73 reveals substitutions at position 3 (tyrosine vs. phenylalanine), potentially introducing additional polar interactions or affecting aromatic stacking at the substrate interface. Similarly, at position 7, *Ec*CBM73 harbours a tyrosine residue, whereas *Vca*CBM73 features a serine, representing a shift from an aromatic to a polar side chain that could influence substrate interactions through altered hydrogen bonding or stacking potential. Interestingly, in *Pa*CBM73, residues corresponding to positions 7 and 8 are entirely absent, resulting in a truncated loop that may contribute to its comparatively narrower binding surface. Such variations, particularly in loop length and residue chemistry, underscore the structural plasticity and potential functional diversification of CBM73s within this clade. Additional substitutions at other positions (e.g., 10 and 12) further emphasize the evolutionary dynamics shaping this domain family.

Together, these structural and sequence-level observations support the idea that CBM73s, particularly those within the same clade, retain a conserved fold but adaptively diverge in surface architecture and residue composition. These variations may allow fine-tuning of binding properties to specific polysaccharide substrates or structural contexts in the broader LPMO scaffold.

### 3.3. EcLPMO is a functional chitinolytic LPMO

In order to probe the activity of *Ec*LPMO, *Ec*LPMO^FL^ and the six truncations thereof were successfully produced in *E. coli*, purified by IMAC, and submitted to several biochemical assays. First, using MALDI-TOF-MS, we verified that *Ec*LPMO^FL^, in the presence of an electron donor (ascorbate), oxidatively cleave α- and β-chitin, as shown by the release of oxidized chitooligosaccharides (CHOS) with degree of polymerization (DP) ranging from DP2 to DP10 (**Fig. S2**). The presence of single and double sodium adducts, [M+Na]^+^ and [M– H+2Na]^+^, is characteristic of C1-oxidized CHOS (aldonic acids), consistent with previous reports (1) (**Fig. S2**). Notably, *Ec*LPMO^FL^ preferentially released even-chain oxidized oligosaccharides from both α- and β-chitin. Monitoring the release of oxidized CHOS revealed that the full-length enzyme *Ec*LPMO^FL^ was more efficient than *Ec*LPMO^CD^ and *Ec*LPMO^ΔCBM73^ over the entire time-course, on both α- and β-chitin (**Fig. S3**). Intriguingly, removal of the CBM73 only nearly abolished the activity, while the *Ec*LPMO^CD^ maintained an intermediate activity. These observations suggest that the DUF domains may play a role in the mode of action of the LPMO on polysaccharides (see more below and in the **Discussion**).

In addition, the Amplex red assay showed that, in the presence of ascorbate, *Ec*LPMO^FL^ reduces O_2_ into H_2_O_2_ (**Fig. S4)**, at a rate comparable to previously characterized LPMOs (15). Experiments with *Ec*LPMO^CD^ and *Ec*LPMO^ΔCBM73^ led to similar values (2-3 x 10^-3^ s^-1^), indicating that, under those conditions, the accessory modules do not affect the reactivity at the copper site (see more in **Discussion**). Of note, control experiments with the *apo* enzyme showed only a low residual activity.

### 3.4. EcCBM73 is crucial for efficient chitin binding

Binding assays were performed with *Ec*LPMO^FL^, its truncated variants (*Ec*LPMO^ΔCBM73^ and *Ec*LPMO^CD^), and the isolated *Ec*CBM73 domain to assess their affinity for α- and β-chitin **(Fig. 3A and B)**. *Ec*CBM73 exhibited the strongest binding, reaching 89.3% and 86.9% binding to α- and β-chitin, respectively, within just 10 minutes **(Fig. 3A and B)**. In comparison, *Ec*LPMO^FL^ showed slower binding kinetics, with maximal binding of approximately 55% to both substrates after 60 minutes **(Fig. 3A and B)**. The truncated variants displayed notably weaker binding: *Ec*LPMO^ΔCBM73^ and *Ec*LPMO^CD^ reached only 38–41% binding to α-chitin (**Fig. 3A**) and 43–57% to β-chitin **(Fig. 3B)**. These results underscore the significant contribution of the CBM73 domain to both the binding kinetics and binding capacity of the full-length enzyme.

**Fig. 3:**
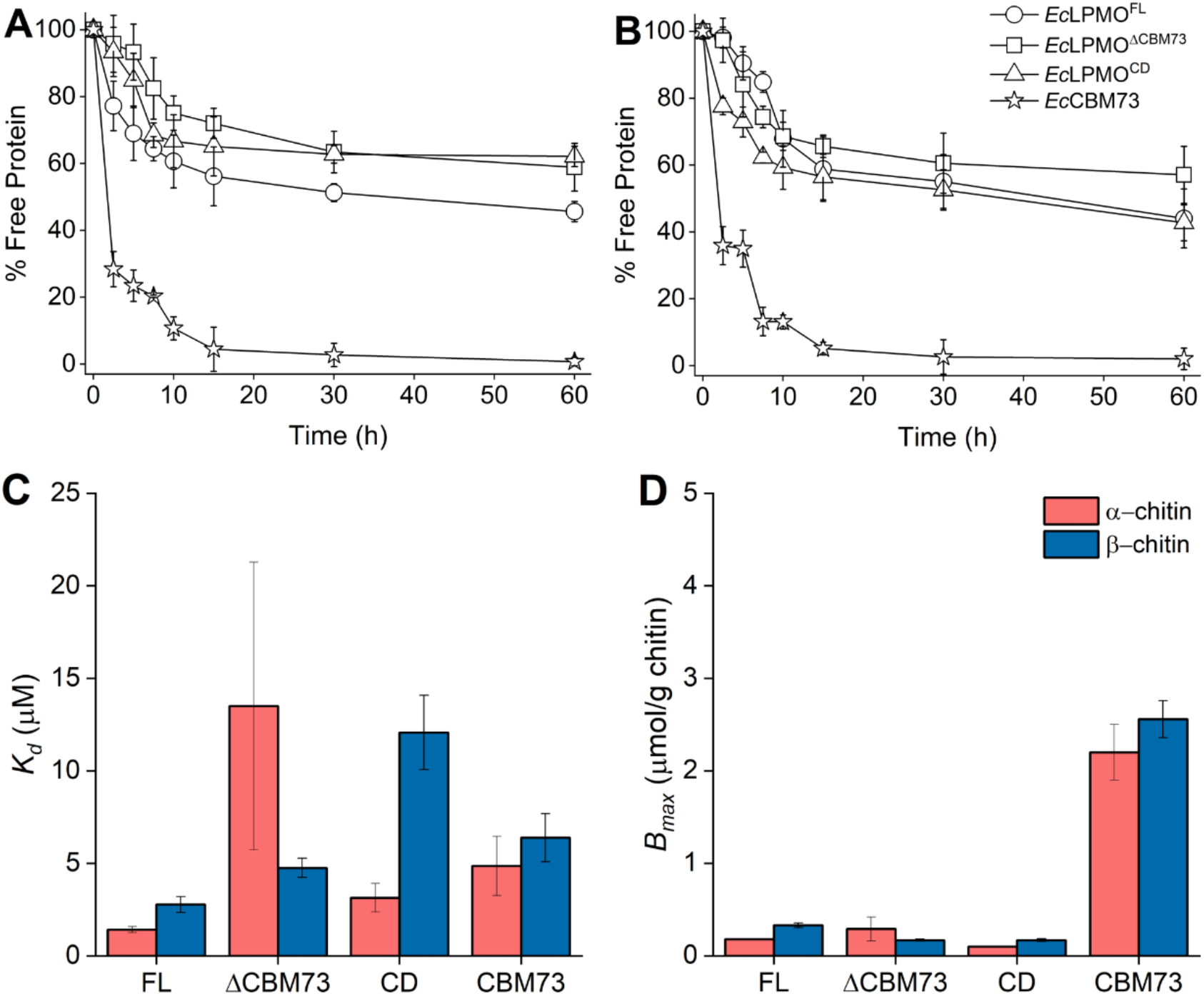
Binding studies of *Ec*LPMO^FL^ and its truncated variants on α- and β-chitin. (A) and (B) Represent the time-course binding of *Ec*LPMO^FL^ and its truncated variants to α- and β-chitin, respectively. The percentage of free protein was determined by measuring the protein concentration in the reaction supernatant collected over time using the BCA method. The experiments were carried out at 22°C using 10 mg/mL substrate in 50 mM sodium phosphate buffer, pH 7.0. (C) and (D) Represent comparison of *K*_d_ (µM) and *B*_max_ (µmol/g chitin), respectively, for *Ec*LPMO^FL^ and its truncated variants, using α- and β-chitin as substrate. Data points (values given in **Table S4**) show average values and error bars show standard deviations (n = 3 independent biological replicates). In panels (C) and (D), *Ec*LPMO^FL^ and its truncated variants, *Ec*LPMO^ΔCBM73^, *Ec*LPMO^CD^ and *Ec*CBM73 have been denoted as FL, ΔCBM73, CD and CBM73, respectively.

To further examine the potential role of other regions of the protein, binding studies were also conducted for the double domain DUF-A+B, and the single domains DUF-A, and DUF-B, on both chitin forms under varying pH conditions. At pH 8.0 and 9.0, none of these domains exhibited appreciable binding to either substrate (**Fig. S5**). However, at pH 10, DUF showed minimal binding (20–24% for both α- and β-chitin; **Fig. S5A, B**), while DUF-A displayed moderate binding, up to 49% to α-chitin and 35% to β-chitin (**Fig. S5C, D**). DUF-B showed no significant binding under any tested condition (**Fig. S5E, F**).

To gain quantitative insights into the binding behaviour, binding isotherms were generated by varying protein concentration at fixed chitin loading, allowing determination of *B*_max_ and *K*_d_ values (**Fig. 3C and D; Table S4**). A clear trend emerged: progressive truncation of *Ec*LPMO^FL^ (*Ec*LPMO^FL^ → *Ec*LPMO^ΔCBM73^ → *Ec*LPMO^CD^) led to increasing divergence in substrate preference between α- and β-chitin. Notably, both *Ec*LPMO^FL^ and *Ec*CBM73 alone showed comparable affinity for α- and β-chitin (based on *K*_d_ and *B*_max_), whereas *Ec*LPMO^ΔCBM73^ and *Ec*LPMO^CD^ exhibited a substantial reduction in affinity (higher *K*_d_) for β - chitin, alongside displaying an expected reduction in *B*_max_ relative to the full-length enzyme (**Fig. 3D; Table S4**). On α-chitin as well, the truncated variants exhibited high *K*_d_ values, indicating reduced substrate affinity, while their *B*_max_ values were comparable to or slightly higher than those of *Ec*LPMO^FL^ (**Fig. 3C; Table S4**).

Indeed, truncation of CBM73 or the removal of additional domains led to a marked loss in substrate affinity, particularly for β-chitin, indicating that these accessory regions modulate substrate preference. Interestingly, despite this reduction in affinity, the truncated variants displayed moderately elevated *B*_max_ values on α-chitin. This apparent paradox, poorer affinity but higher maximal binding, likely reflects increased exposure or conformational flexibility of the catalytic domain in the truncated forms, allowing more enzyme molecules to interact non-specifically or transiently with the substrate. These findings suggest that in the *Ec*LPMO^FL^, the accessory modules, especially CBM73, may restrict the number of catalytically engaged enzymes but confer higher binding specificity and stability. Moreover, the progressive loss of selectivity between α- and β-chitin upon domain deletion implies a possible structural tuning of the full-length enzyme towards substrates that resemble α-chitin more closely. Such domain-mediated modulation of substrate interaction may be critical for function in physiological contexts and highlights the sophisticated role of modularity in fine-tuning LPMO activity.

### 3.5. EcLPMO^FL^ boosts the activity of E. cloacae chitinases

To assess the potential interplay between *E. cloacae* chitinases, specifically *Ec*ChiA, a unimodular GH18 enzyme (53), and *Ec*ChiB, a multimodular enzyme comprising an N-terminal GH18 catalytic domain, a polycystic kidney disease (PKD) domain, and two tandem chitin-binding modules (54), and *Ec*LPMO^FL^, we performed synergy assays using both α- and β-chitin substrates. The addition of *Ec*LPMO^FL^ substantially enhanced the activity of both chitinases on α-chitin (**Fig. 4A, B**) and β-chitin (**Fig. 4C, D**). Notably, *Ec*LPMO^FL^ displayed a particularly strong synergistic effect with *Ec*ChiA, resulting in up to 14-fold and 60-fold increases in GlcNAc release from α- and β-chitin, respectively, compared to reactions containing the chitinase alone (**Fig. 4A, C**). Synergy with *Ec*ChiB was also evident, though less pronounced, yielding up to 2-fold and 1.6-fold increases in (GlcNAc)_2_ release from α- and β-chitin, respectively (**Fig. 4B, D**). Given the pronounced synergistic enhancement observed with *Ec*ChiA, this chitinase was selected for subsequent experiments involving truncated *Ec*LPMO variants.

**Fig. 4:**
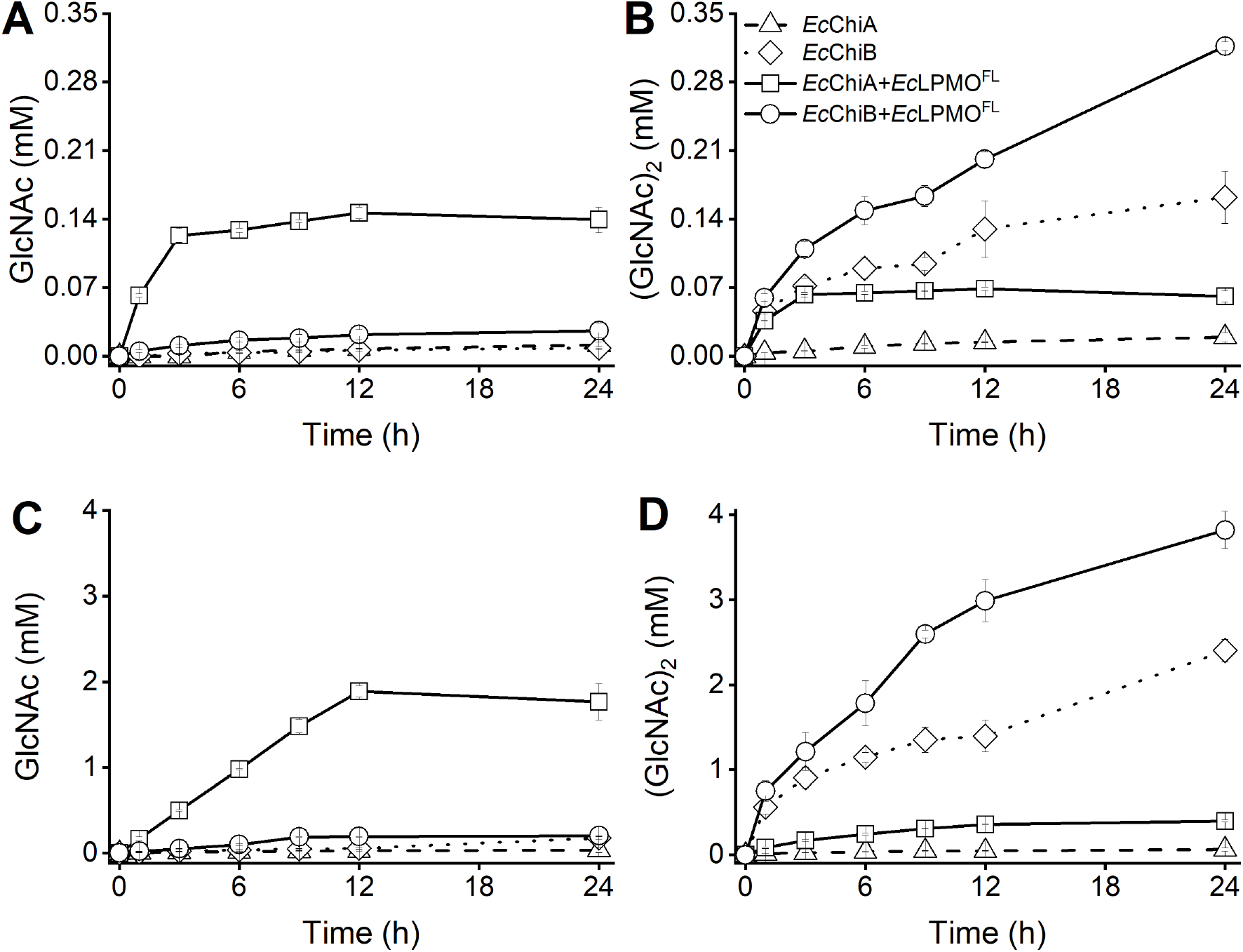
Synergistic chitin degradation by *Ec*LPMO^FL^ and *E. cloacae* chitinases. The graphs show the time-course release of total soluble GlcNAc and (GlcNAc)_2_ by *E. cloacae* chitinases (*Ec*ChiA or *Ec*ChiB; 1 µM), in the absence or presence of *Ec*LPMO^FL^ (1 µM) and ascorbate (1 mM), during the degradation of α-chitin (A and B) and β-chitin (C and D). Data points show average values and error bars show standard deviations (n = 3 independent biological replicates).

### 3.6. Deletion of EcCBM73 impairs the synergistic activity of EcLPMO with chitinases

To evaluate the contribution of *Ec*LPMO^FL^ modular structure to its synergistic function with *Ec*ChiA, we compared the full-length enzyme with its truncated variants, *Ec*LPMO^ΔCBM73^ and *Ec*LPMO^CD^. Overall, in contrast to *Ec*LPMO^FL^, both truncated variants showed markedly reduced performance on both α- and β-chitin (**Fig. 5A**). More in details, on α-chitin, the truncated variants resulted in nearly a two-fold reduction in GlcNAc yield at 24 h compared to the full-length enzyme (**Fig. 5A**). Similarly, (GlcNAc)_2_ yields dropped by ∼2.9-fold and ∼2.6-fold for *Ec*LPMO^ΔCBM73^ and *Ec*LPMO^CD^, respectively (**Fig. 5B**). For β-chitin, GlcNAc release decreased by up to 5.5-fold in reactions with *Ec*LPMO^ΔCBM73^, while no detectable products were observed with *Ec*LPMO^CD^ beyond the 24 h mark (**Fig. 5C**). This trend was mirrored in (GlcNAc)_2_ release profiles (**Fig. 5D**). Together, these findings highlight the essential role of the CBM73 domain in sustaining and enhancing the synergistic action of *Ec*LPMO^FL^ with chitinases.

**Fig. 5:**
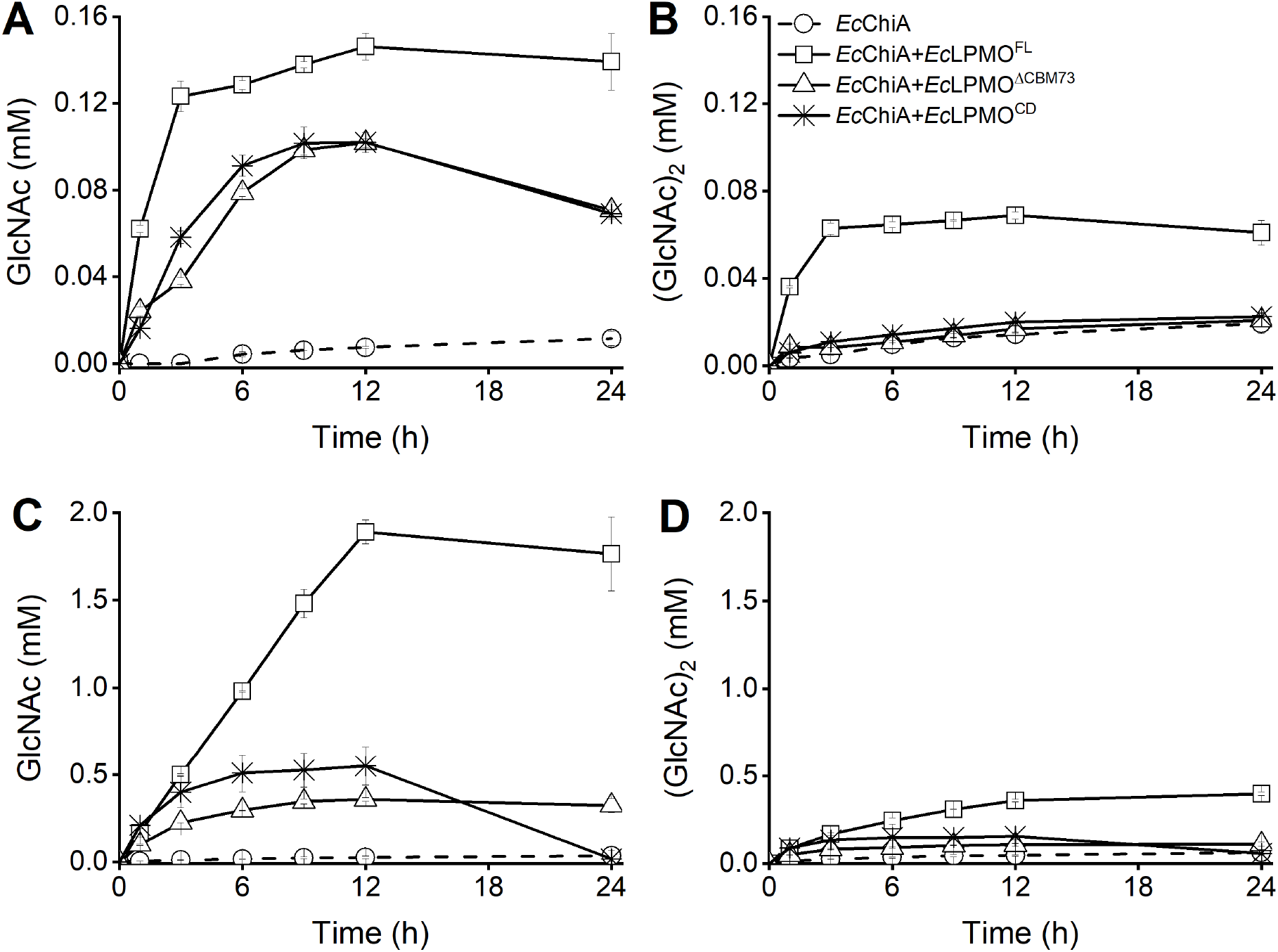
Synergistic effect of *Ec*LPMO^FL^ and its truncated variants on crystalline chitin degradation by *Ec*ChiA. The graphs show the time-course release of total soluble GlcNAc and (GlcNAc)_2_ by *Ec*ChiA (1 µM), in the absence or presence of *Ec*LPMO^FL^ and its truncations (*Ec*LPMO^ΔCBM73^ and *Ec*LPMO^CD^) (1 µM) and ascorbate (1 mM), during the degradation of α-chitin (A and B) and β-chitin (C and D). Data points show average values and error bars show standard deviations (n = 3 independent biological replicates).

### 3.7. The chitinolytic machinery of Enterobacter cloacae

In an attempt to draw a broader picture of *Ec*LPMO function, we examined its genomic context, unveiling an elaborate chitinolytic system comprising two GH18 chitinases, two GH19 chitinases, two *β*-N-acetylhexosaminidases (GH3 and GH20), and one CE4 polysaccharide deacetylase. Several of these enzymes are genomically co-localized, indicating potential functional associations (**Fig. 6**). *Ec*LPMO is located in the same operon as *Ec*HexA (GH20), while *Ec*ChiB and *Ec*ChiC are adjacent to a predicted type II secretion system (**Fig. 6A**). The remaining genes are dispersed. Among the encoded chitin-active CAZymes, *Ec*ChiA, *Ec*ChiB, *Ec*ChiC, *Ec*HexA and *Ec*LPMO, possessed an N-terminal signal peptide, indicating extracellular secretion (**Fig. 6B**). An illustration of chitin degradation mediated using these secreted enzymes by *E. cloacae* have been represented in **Fig. 6C**. This chitinolytic gene repertoire is conserved across *E. cloacae* strains, with *Ec*LPMO among the most conserved components (**Table S2**).

**Fig. 6:**
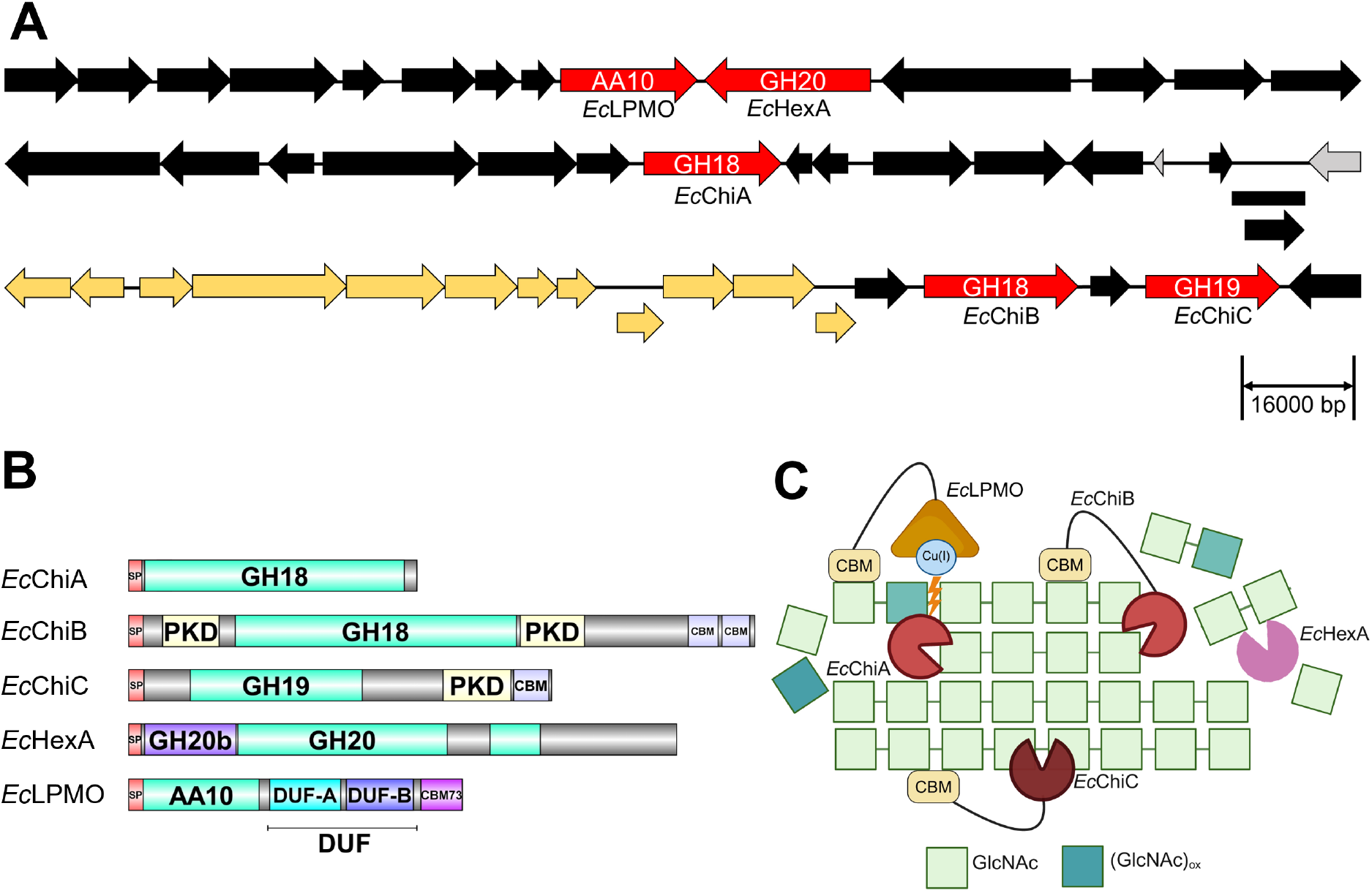
Schematic representation of extracellular chitin degradation by *E. cloacae*. (A) Represents the genetic loci depicting gene arrangement of the chitin-active CAZymes of *E. cloacae* ATCC 13047^T^ that possess an N-terminal signal peptide, indicating extracellular secretion. The direction of the arrows indicates the strand on which they are located. Genes with overlapping loci are indicated with arrows outside the frame. The size of the arrows represents the gene size (scale in base pairs provided at the bottom of the figure). The arrows color code is as follows: red, genes coding for the chitin-active CAZymes (the label within the arrow indicates the CAZyme family, while the one at the bottom represents the name of the gene/protein); yellow, genes involved in the general secretory pathway; grey, genes encoding hypothetical proteins; black, unrelated genes. (B) Represents the domain architecture of these chitin-active CAZymes. Abbreviations: SP – Signal Peptide; GH – Glycoside Hydrolase; Hex-β-N-acetylhexosaminidase; PKD – Polycystic Kidney Disease domain; CBM – Carbohydrate Binding Module; GH20b – N-terminal domain of GH20; DUF – Domain of Unknown Function. (C) Represents an illustration of chitin degradation mediated by *E. cloacae* using the secreted enzymes shown in (A and B). Note: *Ec*ChiC and *Ec*HexA are yet to be characterized and hence their representation was made based on CAZy database prediction and existing literature.

## 4. Discussion

LPMOs are a unique class of enzymes originally identified for their critical role in the oxidative cleavage of polysaccharides during biomass degradation. However, emerging studies have expanded their functional scope (26), revealing involvement in other biological contexts, including their roles as virulence factors in bacterial pathogenesis. These include AA10 LPMOs from human pathogens – viz., the *Lm*LPMO10 from *Listeria monocytogenes* (55, 56), GbpA from *Vibrio cholerae* (32, 57, 58), and CbpD from *Pseudomonas aeruginosa* (27), along with proteins from pathogens affecting animals, such as fusolin from *Entomopoxvirus* (19), *Pl*CBP49 from *Paenibacillus larvae* (59), and *As*LPMO10A/B from *Aliivibrio salmonicida* (60). These findings have broadened the understanding of LPMO function across diverse pathogenic bacteria.

In this study, we examined an AA10 LPMO from an important bacterial human pathogen, namely *E. cloacae*. Interestingly, analysis of the available genomes of *E. cloacae* strains revealed that most strains (40 out of 46) encode a single *lpmo* gene (**Table S1**), suggesting strong conservation and possible functional indispensability of this gene.

*Ec*LPMO is a GbpA-type LPMO with a modular organization resembling GbpA from *V. cholerae* (32), CbpD from *P. aeruginosa* (27), and *Vca*LPMO10A from *V. campbellii* (35). We showed here that *Ec*LPMO, alike the latter LPMOs, can bind and oxidize crystalline chitin. Moreover, it showed notable synergy with the GH18 chitinase *Ec*ChiA, enhancing chitin breakdown to GlcNAc. In contrast, co-incubation with the multi-modular *Ec*ChiB primarily yielded (GlcNAc)_2_, highlighting differential synergy between *Ec*LPMO and chitinases. Diving further into the boosting effect of *Ec*LPMO, we found that the CBM73 plays a crucial role as shown by the reduced chitin degradation capacity of *Ec*LPMO^ΔCBM73^ and *Ec*LPMO^CD^ variants and their lack of synergy with *Ec*ChiA. The strong chitin-binding properties of *Ec*CBM73, similar to its homolog in other bacteria (27, 40) albeit with some differences in binding strength, probably explain its important role in sustaining *Ec*LPMO function. It has indeed been shown that productive polysaccharide binding is key to prevent LPMO oxidative auto-inactivation and thus maximize its boosting potential (11, 61, 62).

Despite several studies, including the present one, on multi-modular LPMOs from pathogenic bacteria, the role of the DUF domains in multimodular LPMOs has remained hitherto mysterious. We have shown here that the DUF-A and DUF-B domains have very limited chitin-binding properties (**Fig. S5**). During the preparation of the present manuscript, the release of AF3 software (46) has allowed to generate new models in the presence of copper, which revealed an unprecedented observation: a unique inter-domain arrangement in which the DUF-B domain folds onto the catalytic AA10 domain, forming an extended metal coordination environment composed of four histidine residues (**Fig. 7, Fig. S6**). In this configuration, two histidines from the AA10 domain coordinate the catalytic copper ion, while two additional histidines from the DUF-B domain complete a predicted tetra-histidine coordination sphere. This predicted inter-domain rearrangement is lost upon mutation of the histidine residues from the DUF-B domain (H354A and H384A in *Ec*LPMO; H357A and H388A in *Vc*GbpA), suggesting that both these residues and Cu(II) are essential for its formation. Notably, confidence in the relative positioning of DUF-A and DUF-B increases markedly when Cu(II) is included during modelling, as reflected in the PAE data for both proteins, further underscoring the potential role of copper in stabilizing the overall tertiary structure (**Fig. 7**).

**Fig. 7:**
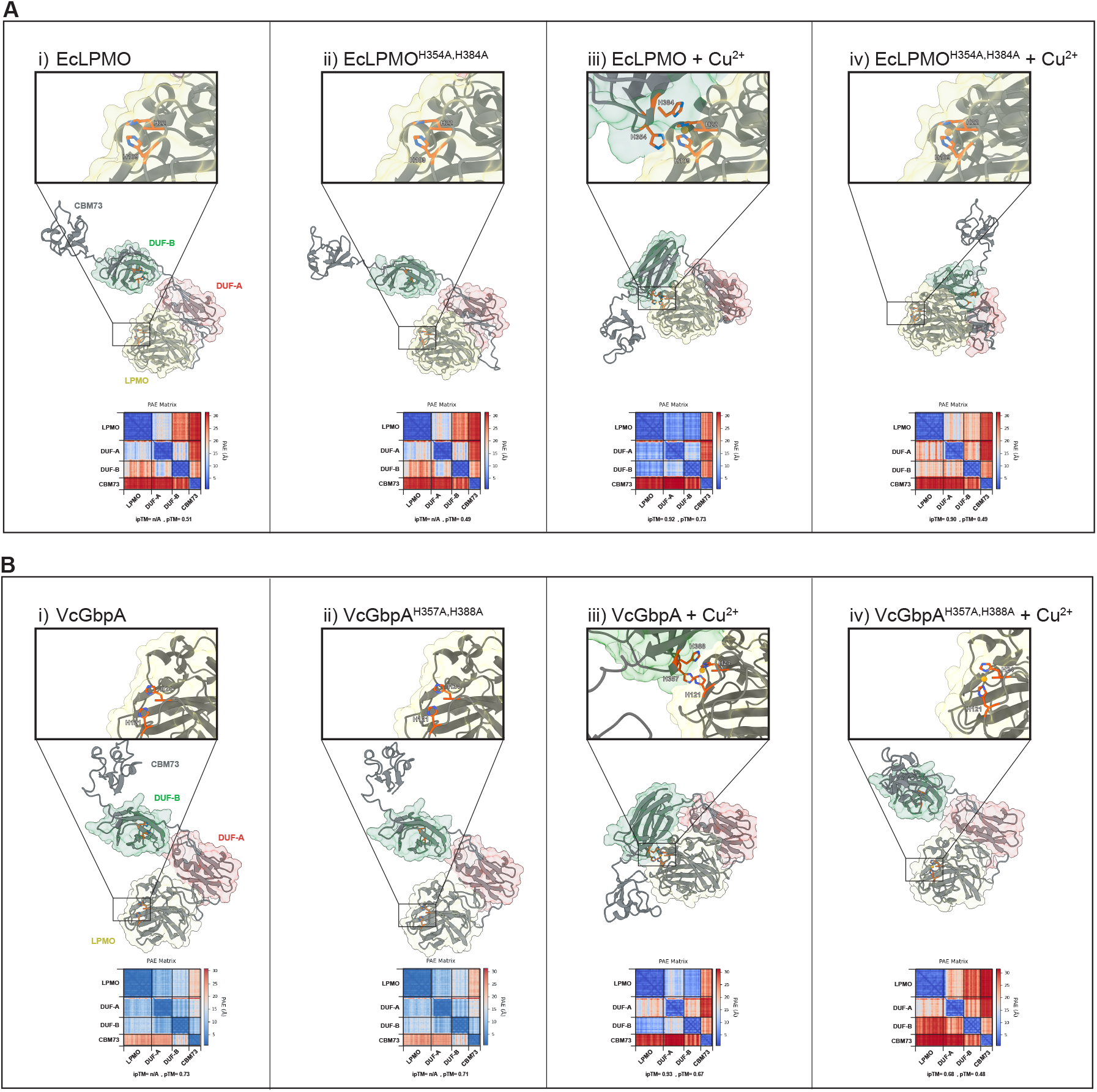
AlphaFold3-predicted structures of mature *Ec*LPMO and *Vc*GbpA including wild-type and His-to-Ala mutants, modelled alone or in complex with Cu(II). (A) (i-iv) Cartoon representations of AlphaFold3 (AF3) models are shown for wild-type mature *Ec*LPMO and the double mutant *H354A, H384A*, each modelled with and without Cu(II). Domains are shaded based on InterProScan annotations: LPMO (yellow), DUF-A (red), and DUF-B (green). Insets highlight the histidine brace region, with histidine side chains rendered as sticks and colored in red. Corresponding Predicted Alignment Error (PAE) plots are shown for each model, alongside the pTM and ipTM confidence metrics. All models were generated using AlphaFold3 with fixed seed number 1. (B) (i-iv) AF3 Models of wild-type mature *Vc*GbpA and the *H357A, H388A* mutant, with and without Cu(II), are shown using the same visual scheme.

The analysis of structural models from other multi-modular LPMOs suggests a conservation of this tetra-histidine copper-coordinated center (**Fig. S6)**. This configuration is, to our knowledge, being reported for the first time and represents a previously unrecognized feature of multi-modular LPMOs. Given the known susceptibility of LPMO active sites to oxidative self-inactivation (11, 61, 63-65), particularly under uncoupled turnover conditions where reactive oxygen species may accumulate, this tetra-histidine arrangement may reflect an evolutionary adaptation to protect the catalytically essential histidine brace from oxidative damage. Thus, in complement to substrate binding (11, 61, 63-65), and second sphere aromatic residues (66, 67), we propose that such unique conformation may represent a new protective strategy. Such configuration could be a way for the microorganism to control the LPMO activity, by for instance rendering the AA10 domain functional upon unfolding of the multi-modular structure only upon binding of its proper substrate. Both hypotheses await, however, to be experimentally tested.

In this context, the strong chitin-binding affinity of CBM73 may serve a dual role beyond substrate recruitment. Based on our results with *Ec*LPMO truncation variants, where *Ec*LPMO^ΔCBM73^ displayed minimal or no activity while *Ec*LPMO^CD^ retained measurable activity, it is tempting to propose that CBM73-mediated substrate engagement facilitates a conformational unlocking of the catalytic site. Specifically, substrate binding may induce structural rearrangement that destabilizes or ‘unshields’ the DUF-B domain from its folded position over the histidine brace, transitioning the enzyme from a protected, inactive state to an open, catalytically competent form. Furthermore, the redox state of the copper might also play an important role in the ‘unshielding’ process. Indeed, the fact that the full-length and truncated enzymes produce H_2_O_2_ at similar rates (**Fig. S4**) suggests that the copper is equally accessible, and, thus, that the tetra-modular copper coordination conformation is not formed under the assay’s reducing conditions. This resonates with previous observations: in its Cu(II) state, the copper ion bound to single AA10 LPMO domains typically coordinates (in addition to the His-Brace) two water molecules (68), which are presumably replaced here by the two DUF-B histidines. Upon reduction of Cu(II) to Cu(I), a coordination site is lost in single LPMO domains (one water molecule is excluded; 68), which may, here, be translated in the disruption of the tetra-histidine coordination, explaining thereby the similar Cu(I) reactivity between WT and truncated enzymes. These possibilities, though speculative, offer compelling directions for future structural and biochemical investigation.

Taking into account all data presented in this study, we propose that the chitinolytic machinery of *E. cloacae* may aid nutrient acquisition in chitin-rich environments such as soil. Alternatively, or in complement, the relevance of these enzymes during host colonization or infection remains to be explored. Indeed, among *Ec*LPMO homologs, GbpA from *V. cholerae* has been studied for its effects on host cells, including mitochondrial dysfunction, ROS production, and NF-κB activation, leading to necrotic cell death (69). In contrast, CbpD from *P. aeruginosa* contributes to immune evasion by inhibiting complement activation and resisting the membrane attack complex (27). Although *Ec*LPMO shares domain architecture with these proteins, its low sequence similarity suggests divergent functions. Nevertheless, its strong conservation across *E. cloacae* strains implies a potentially important role in host-pathogen interactions.

## 5. Conclusion

In summary, *Ec*LPMO represents a distinct multi-modular LPMO with chitin-binding and oxidative activity, functioning in synergy with endogenous chitinases. Importantly, the observations made on the individual roles of each module, and notably the presence of putative inter-domain tetra-histidine copper coordination center, expand our understanding of LPMO structural diversity and suggest potential avenues for engineering LPMOs with enhanced oxidative resilience and stability. Furthermore, its conserved presence across *E. cloacae* strains, along with its modular similarities to known virulence-associated LPMOs, points to a potentially broader biological role, including in host-pathogen interactions. These findings highlight *Ec*LPMO as a promising candidate for further structural and functional studies, both in the context of microbial pathogenesis and environmental chitin turnover.

## Acknowledgements

We are grateful to Prof. Appa Rao Podile (Department of Plant Sciences, School of Life Sciences, University of Hyderabad) for kindly providing the genomic DNA of *E. cloacae* 13047^T^. We also thank Prof. Bernard Henrissat and the CAZy team in Marseille, France, for their support in annotating the domain boundaries, specifically AA10, DUF-A, DUF-B, and CBM73.

