## Supplementary Information for "Functional insights into a peculiar tetra-modular LPMO from the human pathogen *Enterobacter cloacae*"

**Table S1:** Comparative analysis of the number of genes coding for chitin-active CAZymes found in the genome of various strains of *Enterobacter cloacae***.**

| S. No | Strain | Isolation source | NCBI Accession | GH18 | GH19 | GH3 | GH20 | AA10 | CE4 | Reference |
|---|---|---|---|---|---|---|---|---|---|---|
| 1. | 1382 | Hospital (Diseased human) | OW968328.1 | 2 | 1 | 1 | 1 | 1 | 1 | N.a.p. |
| 2. | FDAARGOS 1431 | Clinical sample | CP077211.1 | 2 | 2 | 1 | 1 | 1 | 1 | N.a.p. |
| 3. | ATCC 13047$^T$ | Human Cerebrospinal fluid | CP001918.1 | 2 | 2 | 1 | 1 | 1 | 1 | This study; (1) |
| 4. | CZ862 | Hospital environment | CP073310.1 | 1 | 2 | 1 | 1 | 1 | 1 | (2) |
| 5. | EFN743 | Human urine | CP092635.1 | 2 | 1 | 1 | 1 | 1 | 1 | N.a.p. |
| 6. | PIMB10EC27 | Human urine | CP020089.1 | 2 | 1 | 1 | 1 | 1 | 1 | N.a.p. |
| 7. | WP5-S18-CRE-02 | Wastewater | AP022126.1 | 2 | 2 | 1 | 1 | 1 | 1 | (3-5) |
| 8. | Isolate-F | Human | CP135475.1 | 2 | 1 | 1 | 1 | 1 | 1 | N.a.p. |
| 9. | MY490 | Chicken | CP053568.1 | 2 | 3 | 1 | 1 | 1 | 1 | N.a.p. |
| 10. | CBG15936 | Human sputum | CP046116.1 | 1 | 2 | 1 | 1 | 1 | 1 | N.a.p. |
| 11. | TL15 | Water | CP126341.1 | 2 | 1 | 1 | 1 | 1 | 1 | N.a.p. |
| 12. | 14240244101 | Human bone | CP045487.1 | 2 | 1 | 1 | 1 | 2 | 1 | N.a.p. |
| 13. | POL9 | Water | CP086408.1 | 2 | 2 | 1 | 1 | 1 | 0 | N.a.p. |
| 14. | AVS0889 | River water | CP092042.1 | 2 | 2 | 1 | 1 | 1 | 1 | (6) |
| 15. | 3143 | Human blood | CP103611.1 | 2 | 3 | 1 | 1 | 1 | 1 | (7) |
| 16. | 2022CK-00409 | Human | CP104836.1 | 1 | 4 | 1 | 1 | 1 | 1 | N.a.p. |
| 17. | M12X01451 | Human stool | CP017475.1 | 1 | 1 | 1 | 1 | 1 | 1 | N.a.p. |
| 18. | ECl_1.VN | Rectal swab of pregnant women | CP085080.1 | 2 | 1 | 1 | 1 | 1 | 1 | N.a.p. |
| 19. | STN0717-73 | Hospital sewage tank | AP022519.1 | 1 | 2 | 1 | 1 | 0 | 1 | N.a.p. |
| 20. | RMCH-M23-N | *Blattella germanica* | CP123598.1 | 2 | 1 | 1 | 1 | 1 | 1 | N.a.p. |
| 21. | Isolate-B | Human | CP135498.1 | 4 | 1 | 1 | 1 | 1 | 1 | N.a.p. |
| 22. | 99183042147 | Human rectum | CP134674.1 | 2 | 1 | 1 | 1 | 1 | 1 | N.a.p. |
| 23. | GGT036 | Forest soil | CP009756.1 | 2 | 1 | 1 | 1 | 1 | 1 | (8) |
| 24. | D41-sc-1712200 | Unknown | CP056779.1 | 2 | 2 | 1 | 1 | 1 | 1 | N.a.p. |
| 25. | EN3600 | Human blood | CP035633.1 | 1 | 3 | 1 | 1 | 0 | 1 | N.a.p. |
| 26. | GX1Z-1L | Swine | CP071861.1 | 1 | 2 | 1 | 1 | 1 | 1 | N.a.p. |
| 27. | CZ-1 | Paddy soil | CP035738.1 | 3 | 3 | 1 | 1 | 3 | 1 | N.a.p. |

| | | | | | | | | | | |
|---|---|---|---|---|---|---|---|---|---|---|
| 28. | SD21 | Human | CP093914.1 | 2 | 1 | 1 | 1 | 1 | 1 | N.a.p. |
| 29. | WP5-S18-ESBL-01 | Wastewater | AP022133.1 | 2 | 1 | 1 | 1 | 0 | 1 | (3-5) |
| 30. | Ecc8276_LB-HALD | Human urine | CP130342.1 | 2 | 1 | 1 | 1 | 1 | 1 | N.a.p. |
| 31. | 12961-yvys | Human urine | CP083821.1 | 2 | 0 | 1 | 1 | 1 | 1 | (9) |
| 32. | SDM | Soil | CP003678.1 | 2 | 1 | 1 | 1 | 1 | 1 | (10) |
| 33. | ECNIH2 | Sink drain | CP008823.1 | 1 | 2 | 1 | 1 | 1 | 1 | (11) |
| 34. | Effluent_3 | Treated sewage effluent | CP039311.1 | 2 | 3 | 1 | 1 | 1 | 1 | N.a.p. |
| 35. | Effluent_4 | Treated sewage effluent | CP039303.1 | 2 | 3 | 1 | 1 | 1 | 1 | N.a.p. |
| 36. | Effluent_2 | Treated sewage effluent | CP039318.1 | 2 | 3 | 1 | 1 | 1 | 1 | N.a.p. |
| 37. | DSM 2648 | Dept. Medical Microbiol, Stanford Univ. | CP056117.1 | 2 | 2 | 0 | 1 | 1 | 1 | N.a.p. |
| 38. | MBRL 1077 | Human wound | CP014280.1 | 4 | 2 | 1 | 1 | 2 | 1 | (12) |
| 39. | Colony340 | Food | CP078540.1 | 2 | 1 | 1 | 1 | 1 | 1 | N.a.p. |
| 40. | Colony350 | Food | CP078539.1 | 2 | 1 | 1 | 1 | 1 | 1 | N.a.p. |
| 41. | Colony466 | Food | CP078538.1 | 1 | 1 | 1 | 1 | 1 | 1 | N.a.p. |
| 42. | Colony146 | Rectal swab | CP078537.1 | 1 | 1 | 1 | 1 | 1 | 1 | N.a.p. |
| 43. | Colony416 | Food | CP078536.1 | 1 | 1 | 1 | 1 | 1 | 1 | N.a.p. |
| 44. | Colony187 | Swab | CP081292.1 | 2 | 1 | 1 | 1 | 1 | 1 | N.a.p. |
| 45. | SBP-8 | Rhizosphere | CP016906.1 | 2 | 1 | 1 | 1 | 1 | 1 | N.a.p. |
| 46. | NH77 | Human host; Hospital sample | CP040827.1 | 2 | 1 | 0 | 1 | 1 | 1 | N.a.p. |

N.a.p: No associated publication.

**Table S2.** Percentage of strains (out of 46 strains) encoding n members of each CAZy family. For instance, 100% of strains encode strictly 1 member of GH20.

| | | GH3 | GH18 | GH19 | GH20 | AA10 | CE4 |
|---|---|---|---|---|---|---|---|
| Number (n) of CAZy family members | 0 | 4 | 0 | 2 | 0 | 7 | 2 |
| | 1 | 96 | 24 | 52 | 100 | 87 | 98 |
| | 2 | 0 | 70 | 28 | 0 | 4 | 0 |
| | 3 | 0 | 2 | 15 | 0 | 2 | 0 |
| | 4 | 0 | 4 | 2 | 0 | 0 | 0 |

**Table S3.** Primers used in this study.

| Name of the gene | Template used for PCR | Primer combination used | Primer sequence (5'- 3') |
|---|---|---|---|
| *Ec*LPMO$^{FL}$ | *Genomic DNA* | *Ec*LPMO-FL-Fp | AATAACATATGAAATTATCAAAAATTGCC |
| | | *Ec*LPMO-FL-Rp | AATAACTCGAGGTGCTTAAGCACCCACGC |
| *Ec*LPMO$^{\Delta CBM73}$ | *Ec*LPMO$^{FL}$ | *Ec*LPMOΔCBM73-Fp | AATAACATATGAAATTATCAAAAATTGCC |
| | | *Ec*LPMOΔCBM73-Rp | AATAACTCGAGGTCCGCCGCGGCTTCCTC |
| *Ec*LPMO$^{CD}$ | *Ec*LPMO$^{FL}$ | *Ec*LPMO-CD-Fp | AATAACATATGAAATTATCAAAAATTGCC |
| | | *Ec*LPMO-CD-Rp | AATAACTCGAGGCCGCCGTTACCAAAATC |
| *Ec*X197 | *Ec*LPMO$^{FL}$ | X197-Fp | AATAACATATGGGTGCCGGCATCGAATCA |
| | | X197-Rp | AATAACTCGAGGTCCGCCGCGGCTTCCTC |
| *Ec*X427 | *Ec*LPMO$^{FL}$ | X427-Fp | AATAACATATGGGTGCCGGCATCGAATCA |
| | | X427-Rp | AATAACTCGAGCTCTTCGTACGAAATCGCAAC |
| *Ec*X428 | *Ec*LPMO$^{FL}$ | X428-Fp | AATAACATATGAATGAAACCATCGCCGTCAGC |
| | | X428-Rp | AATAACTCGAGGTCCGCCGCGGCTTCCTC |
| *Ec*CBM73 | *Ec*LPMO$^{FL}$ | CBM73-Fp | AATAACATATGCATGATTACATCTTCCCG |
| | | CBM73-Rp | AATAATCTAGAGTTGCAGGTGCCCAGATC |

**Table S4:** Binding kinetic parameters, $K_d$ (μM) and $B_{max}$ (μmol/g chitin) values for *Ec*LPMO$^{FL}$ and its truncated variants, using α- and β-chitin as substrate.

| | **α-chitin** | | **β-chitin** | |
|---|---|---|---|---|
| **Proteins** | $K_d$ | $B_{max}$ | $K_d$ | $B_{max}$ |
| *Ec*LPMO$^{FL}$ | 1.71±0.49 | 0.82±0.15 | 2.73±0.51 | 0.69±0.08 |
| *Ec*LPMO$^{\Delta CBM73}$ | 13.5±7.78 | 0.29±0.13 | 4.75±0.52 | 0.17±0.011 |
| *Ec*LPMO$^{CD}$ | 3.14±0.77 | 0.1±0.009 | 12.07±2.01 | 0.17±0.016 |
| *Ec*CBM73 | 4.86 ± 1.6 | 2.2 ± 0.3 | 6.38 ± 1.3 | 2.56 ± 0.2 |

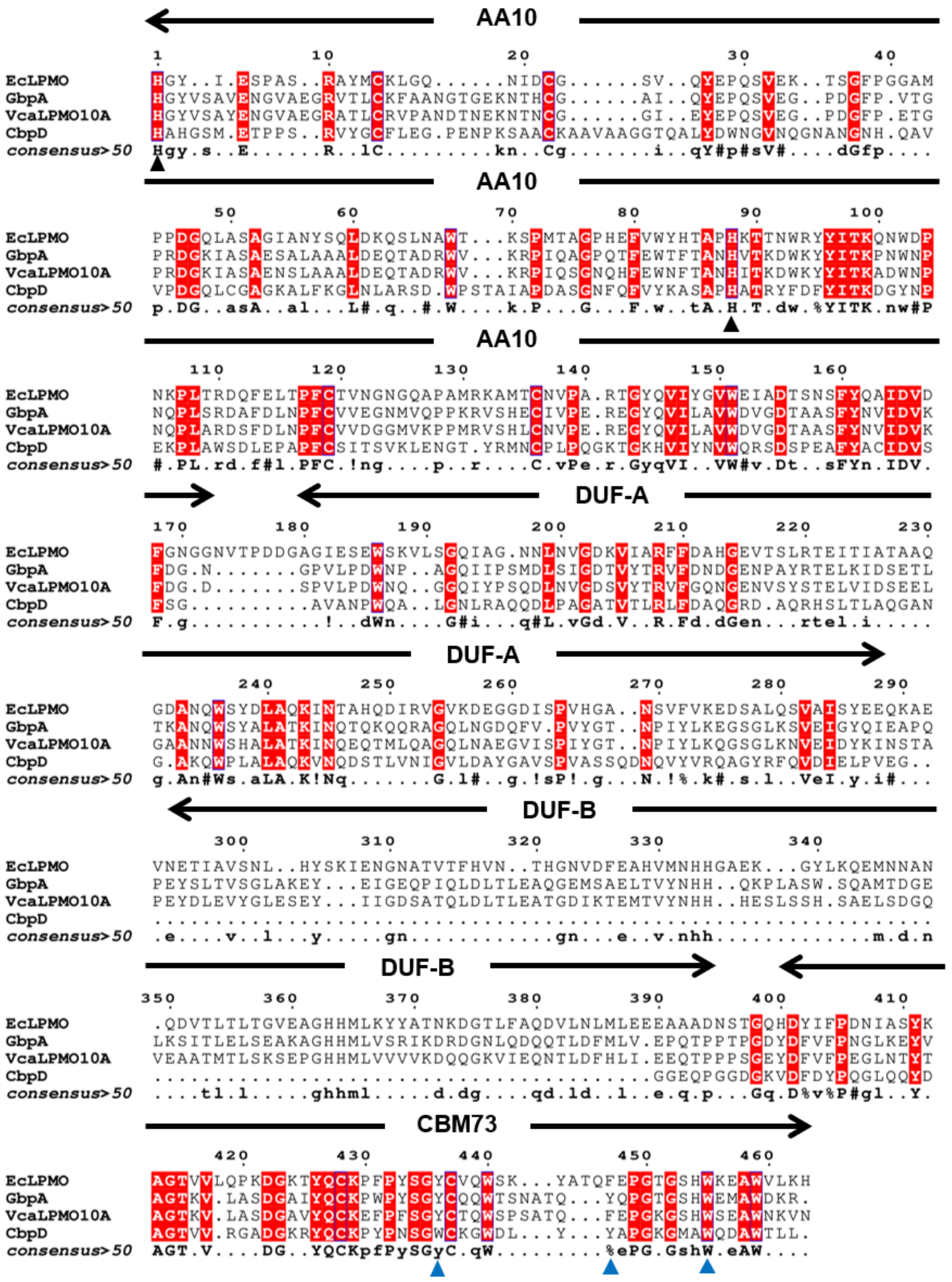

**Fig. S1: Multiple sequence alignment of *Ec*LPMO against GbpA, *Vca*LPMO10A, and CbpD.** Conserved residues are shown in red. The multiple sequence alignment was performed using T-Coffee Simple MSA online server and the output was generated using ESPript 3.0. The domain demarcation in the MSA was done according to *Ec*LPMO. The conserved histidine residues of the histidine brace in the AA10 domain are indicated with black triangles at positions His1 and His88 of the MSA. The residues lining the chitin-binding surface in CBM73 [as per Madland et al., 2021(13)] are indicated with blue triangles.

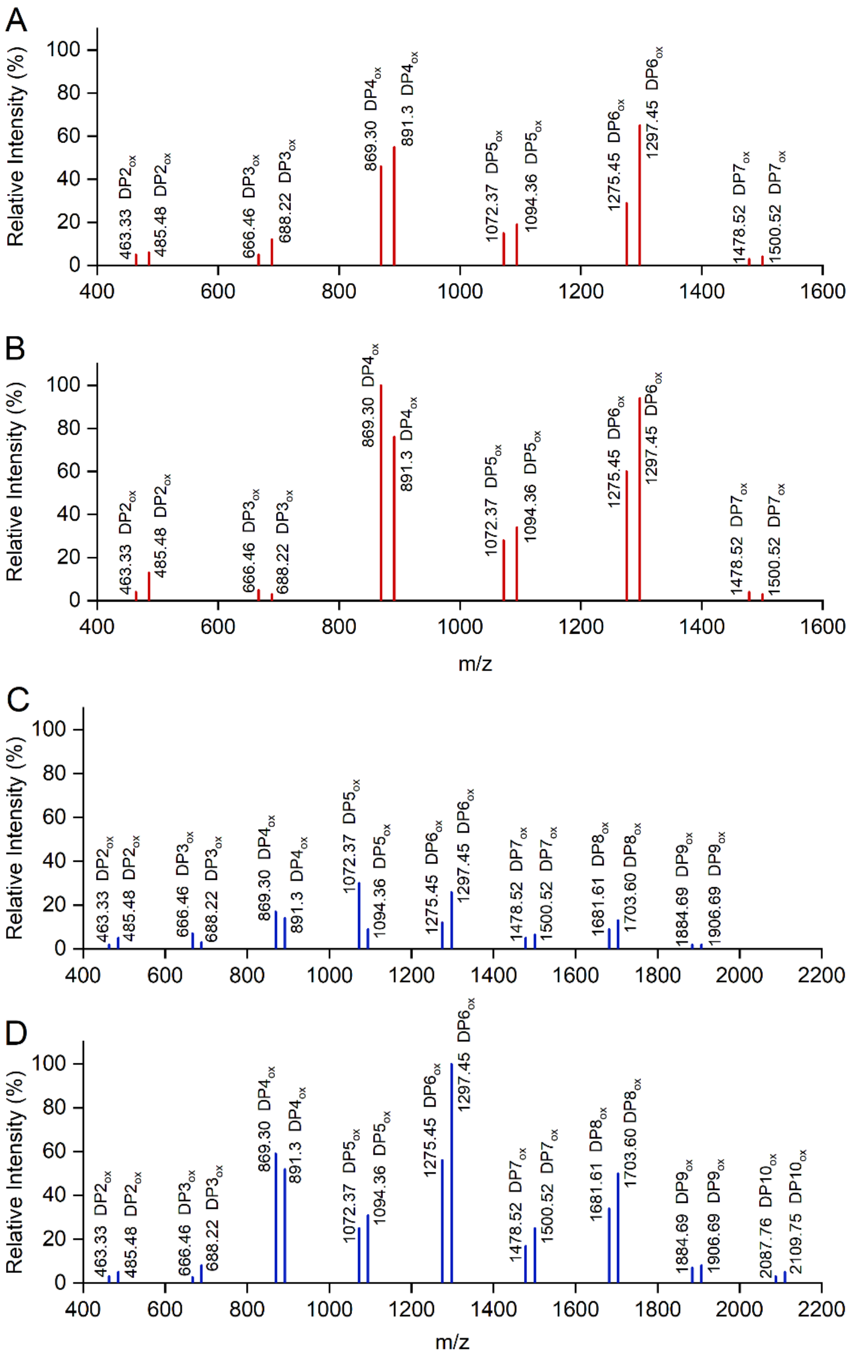


**Fig. S2: Mass spectrometry analysis of *Ec*LPMO products.** The figure shows the distribution of m/z values of oxidized chitooligosaccharides with varying degrees of polymerization (DP), generated by 1 µM *Ec*LPMO$^{FL}$ acting on 10 mg/mL α-chitin (A & B) and β-chitin (C & D) for 12 h (A & C) and 24 h (B & D). The reactions were conducted in the presence of 1.0 mM ascorbic acid in 5 mM Tris-Cl, pH 8.0, at 37ºC and 1000 rpm. In the MALDI-TOF MS spectra presented here, 100% relative intensity corresponds to $3.8\times10^4$ and $5.1 \times 10^4$ arbitrary units (a.u.) for α- and β-chitin, respectively. For each DP, the two main peaks correspond to the single and double sodium adduct of the aldonic acid, respectively labelled as $[Ald + Na]^+$ and $[Ald\text{-}H\text{+}2Na]^+$.

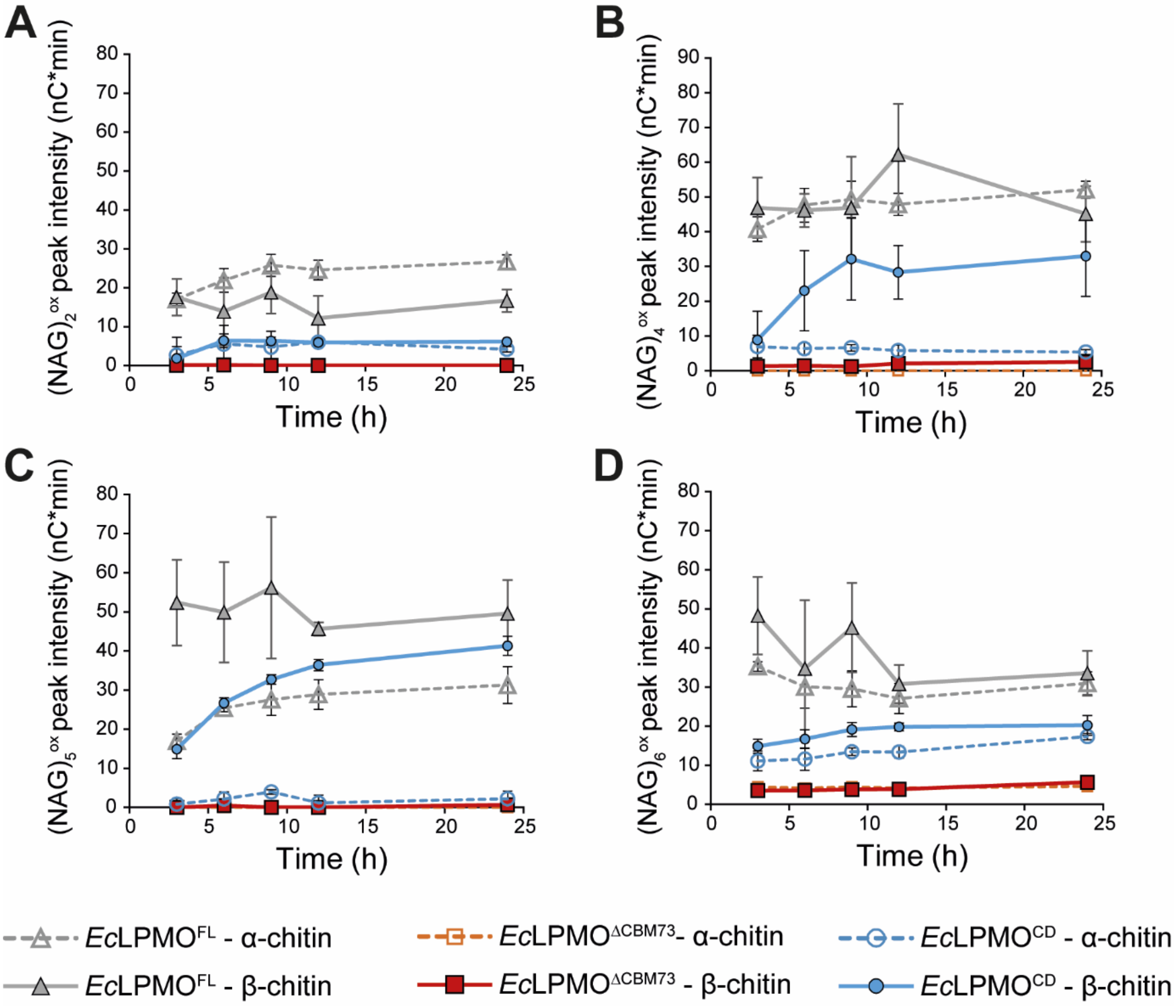


**Fig. S3: Chitinolytic activity of *Ec*LPMO full-length and truncations thereof.** The graphs show the time-course release of C1-oxidized chitooligosaccharides of $(GlcNAc)_2$ (A), $(GlcNAc)_4$ (B), $(GlcNAc)_5$ (C) and $(GlcNAc)_6$ (D) from α- and β-chitin by *Ec*LPMO$^{FL}$, *Ec*LPMO$^{ΔCBM73}$ and *Ec*LPMO$^{CD}$ (the legend provided at the bottom of the figure applies to all panels). All reactions were performed using 1 μM enzyme in the presence of 1.0 mM ascorbic acid in 50 mM Tris-Cl, pH 8.0, at 37°C, 1,000 rpm. Data points show average values and error bars show standard deviations (n = 3 independent biological replicates).

.

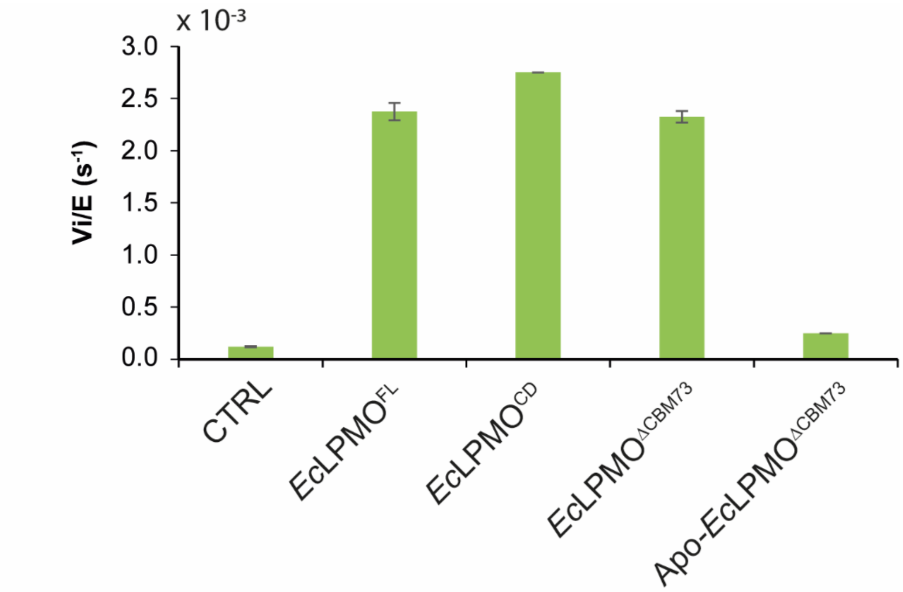


**Fig. S4: *Ec*LPMO $H_2O_2$ production rate.** The graph shows the normalized rate of $O_2$ reduction into $H_2O_2$ by *Ec*LPMO full length protein and truncated variants (1 µM), measured with the Amplex red assay. Two control reactions were carried out: (i) a reaction in the absence of LPMO ("CTRL") showing the non-enzymatic background signal, and (ii) a reaction in which the copper atom has been removed from the LPMO active site ("Apo"). Reactions were carried out in the absence of substrate and in the presence of AscA (50 µM), in sodium phosphate buffer (50 mM, pH 7.0). Data points show average values and error bars show standard deviations (n = 3 independent biological replicates).

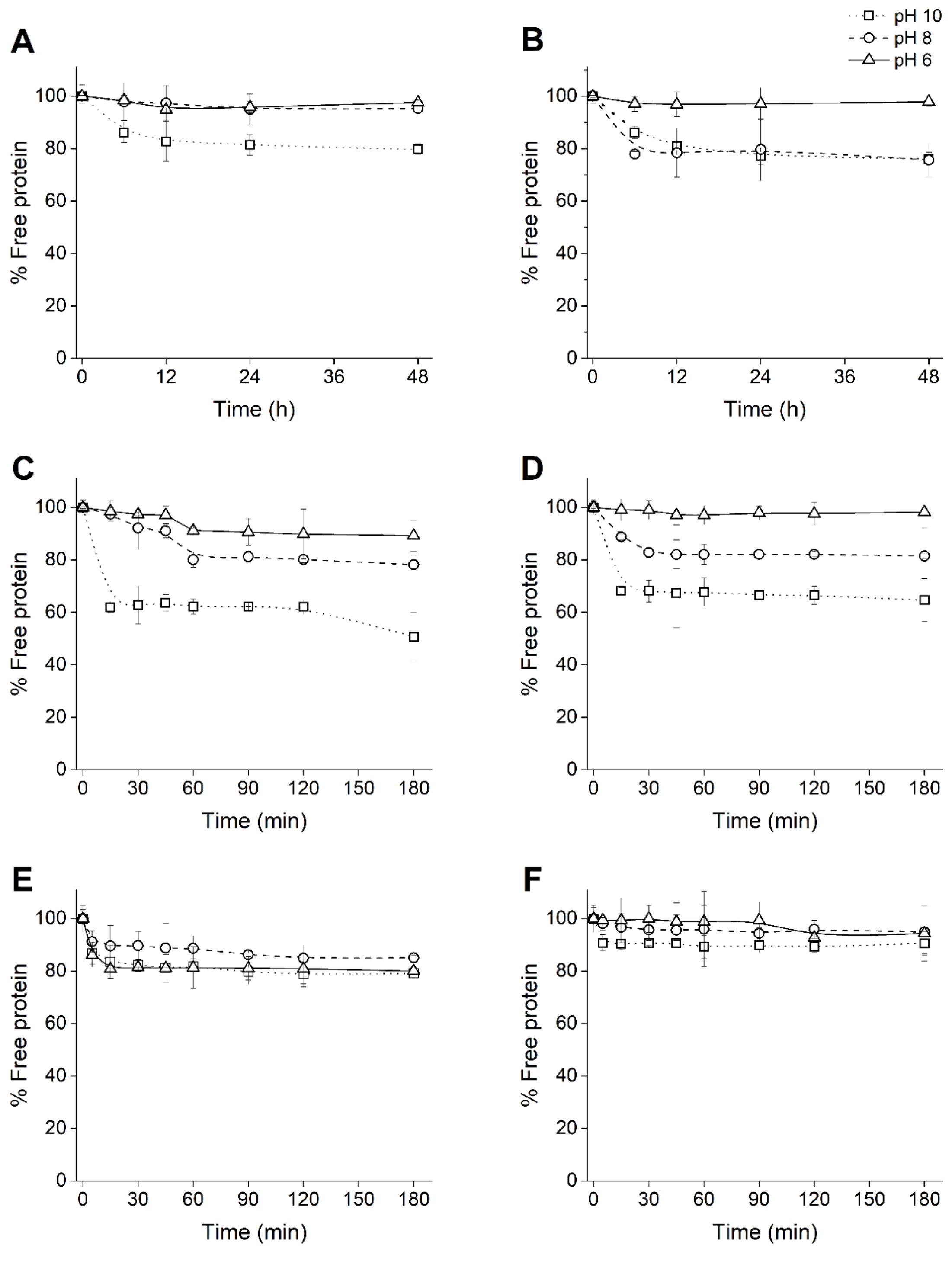


**Fig. S5: Binding of *Ec*LPMO DUF domains (DUF-A+B, DUF-A and DUF-B) to α- and β-chitin.** The graphs show the time course binding of DUF-A+B), DUF-A (C and D), and DUF-B (E and F) to α-chitin (A, C, and E) and β-chitin (B, D, and F) at different pH conditions. Note the different time scales of binding kinetics between the different modules. The percentage of free protein was determined by estimating the protein concentration in the supernatant of the reaction mixtures using the BCA method. The experiments were conducted using 10 mg/mL of α- and β-chitin at 22°C under various pH conditions: 50 mM Bis-Tris buffer (pH 6.0), 50 mM sodium phosphate buffer (pH 8.0) and 50 mM CAPS buffer (pH 10.0). Data points show average values and error bars show standard deviations (n = 3 independent biological replicates).

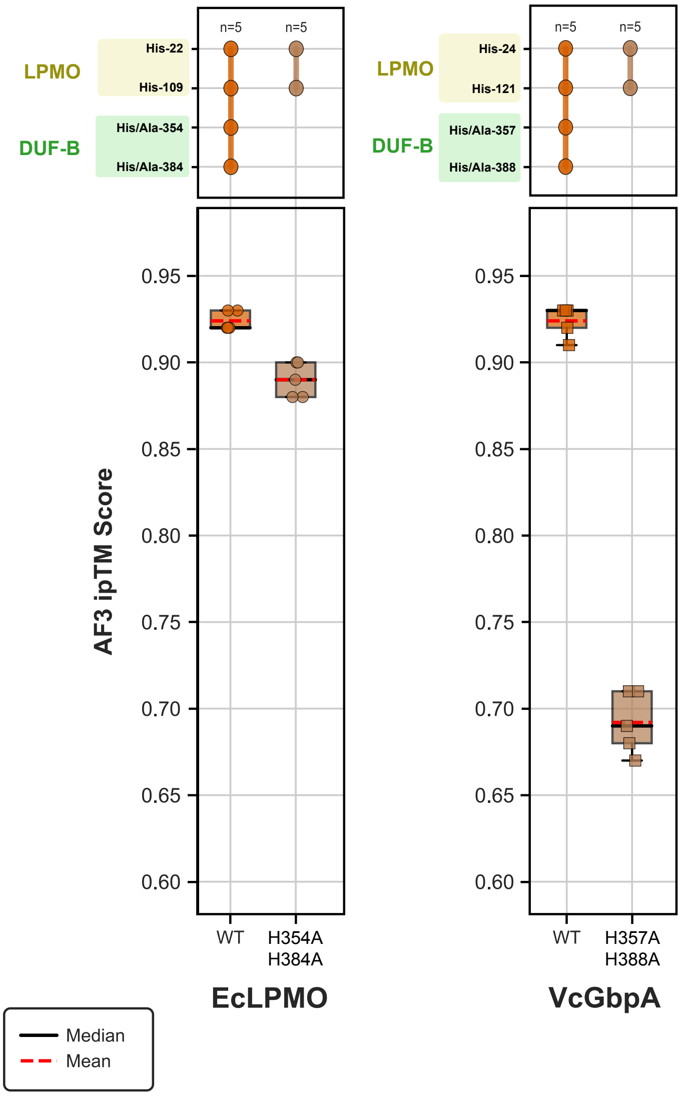


**Fig. S6: Predicted Cu(II) coordination and model confidence scores for *Ec*LPMO and *Vc*GbpA AlphaFold3 models.** AlphaFold3 models were generated using five independent seed values (1–5) for wild-type and double histidine-to-alanine mutants of mature *Ec*LPMO (H354A, H384A) and *Vc*GbpA (H357A, H388A), all in the presence of Cu(II). The ***top panel*** displays an UpSet plot summarizing predicted Cu(II) coordination contacts within a 3.6 Å threshold, specifically with conserved histidine residues in the LPMO and DUF-B domains; n values indicate the number of top-ranked models from each seed, in which each histidine contact pattern was observed. The ***bottom panel*** shows box plots of predicted ion-protein interface confidence scores (ipTM) across the five AlphaFold3 seeds, for every modelled condition.
